# A proteome-wide quantitative platform for nanoscale spatially resolved extraction of membrane proteins into native nanodiscs

**DOI:** 10.1101/2024.02.10.579775

**Authors:** Caroline Brown, Snehasish Ghosh, Rachel McAllister, Mukesh Kumar, Gerard Walker, Eric Sun, Talat Aman, Aniruddha Panda, Shailesh Kumar, Wenxue Li, Jeff Coleman, Yansheng Liu, James E Rothman, Moitrayee Bhattacharyya, Kallol Gupta

## Abstract

The intricate molecular environment of the native membrane profoundly influences every aspect of membrane protein (MP) biology. Despite this, the most prevalent method of studying MPs uses detergent-like molecules that disrupt and remove this vital local membrane context. This severely impedes our ability to quantitatively decipher the local molecular context and comprehend its regulatory role in the structure, function, and biogenesis of MPs. Using a library of membrane-active polymers we have developed a platform for the high-throughput analysis of the membrane proteome. The platform enables near-complete spatially resolved extraction of target MPs directly from their endogenous membranes into native nanodiscs that maintain the local membrane context. We accompany this advancement with an open-access database that quantifies the polymer-specific extraction variability for 2065 unique mammalian MPs and provides the most optimized condition for each of them. Our method enables rapid and near-complete extraction and purification of target MPs directly from their endogenous organellar membranes at physiological expression levels while maintaining the nanoscale local membrane environment. Going beyond the plasma membrane proteome, our platform enables extraction from any target organellar membrane including the endoplasmic reticulum, mitochondria, lysosome, Golgi, and even transient organelles such as the autophagosome. To further validate this platform, we took several independent MPs and demonstrated how our resource can enable rapid extraction and purification of target MPs from different organellar membranes with high efficiency and purity. Further, taking two synaptic vesicle MPs, we show how the database can be extended to capture multiprotein complexes between overexpressed MPs. We expect these publicly available resources to empower researchers across disciplines to efficiently capture membrane ‘nano-scoops’ containing a target MP and interface with structural, functional, and other bioanalytical approaches. We demonstrate an example of this by combining our extraction platform with single-molecule TIRF imaging to demonstrate how it can enable rapid determination of homo-oligomeric states of target MPs in native cell membranes.

## Introduction

The local membrane environment plays a pivotal role in governing every facet of membrane protein (MP) biology. This ranges from the trafficking of MPs to target physiological membranes, formation of various homo and heteromeric signaling complexes, regulation of functional conformational states, to ultimately, degradation of MPs^1,2,3^. Mounting evidence demonstrates that the local membrane intricately controls each aspect of MP life^4,5^. Yet, paradoxically, the most predominant protocol for studying MPs involves the use of micellar detergents, which effectively strip away this crucial local membrane context^6–8^. Recent efforts have turned to polymer-based approaches, such as styrene-maleic acid copolymers (SMA) as an alternative, aiming to preserve the native membrane environment by encapsulating MPs into native nanodiscs^9,10^. These polymer-derived native nanodiscs (SMALP when SMA is the polymer) offer the capability to capture MPs directly from their physiological environment while preserving their native membrane context, presenting a unique opportunity to study membrane biology. The potential application encompasses the ability to discern the conformation dynamics and organization of MPs within the membrane^11,12^, analyze the local membrane environment^13^, and decipher how the intricate interplay between these factors influences cellular signaling^14^. However, establishing these polymers as a widely applicable and versatile toolkit for spatially-resolved membrane biology demands the ability to efficiently capture any target MPs directly from their endogenous membrane, along with their native membrane context, at physiological expression levels. Unfortunately, compared to detergents, these polymers have consistently exhibited markedly lower efficiencies in extracting target MPs. This in turn has severely constrained their application in elucidating membrane-associated cell signaling. This limitation has been consistently noted, including in large-scale comparisons conducted by a major commercial supplier of the polymer, where standard detergents outperformed these polymers^15^. Even our own initial experiments, using overexpressed GFP-tagged AqpZ, yielded considerably lower quantities of extracted protein in polymers compared to commonly used detergents (Extended Data Fig. 1). As a result, these polymer-based approaches have largely been confined to cryoEM studies of MPs, where the target MPs are significantly overexpressed to compensate for the low recovery^16,17^. Even within that, since the introduction of SMA, over the last 15 years, only 5 unique eukaryotic MP structures have been determined using SMA or related polymers (SI Table 1). At the time of writing this manuscript, among the last 150 MP entries in PDB, only 1 has been obtained in SMA or related polymers (Extended Data Fig. 1). This highlights the current extremely limited utility of these native nanodisc-forming polymers; effectively serving as a last resort for cryoEM structure determination of overexpressed MPs where detergents or reconstituted membrane scaffold protein (MSP)-nanodiscs fail to deliver. This fundamentally undermines the remarkable potential that these molecules hold in advancing our understanding of membrane biology by capturing membrane snapshots with nanoscale spatial resolution.

To bridge this technological gap, first, it is imperative to develop protocols and pipelines that enable spatially-resolved extractions of target MPs directly from their native organellar membranes with high efficiency. Such efforts would expand the reach of the technology beyond the abundant MPs and enable applications on low-abundant MPs, directly from their native organelle membranes at their endogenous low expression levels. Secondly, it is essential to have proteome-scale high-throughput resources and platforms that can rapidly deliver the most optimized extraction efficiency for target MPs, as the existing protocols are extremely low throughput. Moreover, the proliferation of various polymers claiming to form native nanodiscs, with many commercially available and others proposed as suitable candidates, adds complexity to the process of identifying the most appropriate chemical conditions for a target MP extraction^18–22^. This makes the entire process both very low throughput and costly. In essence, democratizing a polymer extraction platform as a universally applicable technology for studying spatially-resolved MP-signaling necessitates the development of a proteome-wide, high-throughput approach that provides a quantitative guide for rapid, efficient, and spatially-resolved extraction of target MPs along with their nanoscale local signaling environments. Such a platform will also simultaneously offer general guidelines and benchmarks for assessing the ability of newly synthesized polymers to extract MPs into native nanodiscs, within a biologically relevant context.

Addressing this, here we present a high-throughput, quantitative platform that delivers the most optimized, spatially resolved extraction conditions for 2065 unique membrane proteins, directly from their endogenous membrane, into native nanodiscs. We first developed high-throughput fluorescence-based assays that can be performed in most laboratory setups to evaluate the broadscale native nanodisc forming capabilities of target polymers in a cell type-specific manner. Next, using quantitative proteomics, we have built a proteome-wide database that captures the variable extraction efficiency of 2065 independent MPs across 11 different polymer conditions and provides the most efficient extraction conditions for each. Our database consists of both integral (both polytopic, and bi-topic MPs), and monotopic peripherally associated MPs (together termed as MP from herein) and covers 11 different polymer extraction conditions. We have developed protocols wherein the extraction efficiency of most MPs into membrane active polymers (MAPs) supersedes detergents, enabling studies at endogenous levels of expression. This also extends to all organellar membranes, enabling MP-extraction from any target organellar lipid-bilayer membranes, such as ER, mitochondria (both inner and outer), lysosome, Golgi, and even transient organelles such autophagosomes. To facilitate widespread access to this resource, we have also developed an Open-access web app (https://polymerscreen.yale.edu) through which all this information can be accessed in both individual or multi-protein-specific manner to rapidly obtain the most optimized extraction condition for any target MP, or multi-MP complex. We subsequently validated the capability of such a database through the near-complete extraction of several target MPs directly from their different host organellar membranes. Coupling this with the recently developed native-nanobleach platform^23^, we further demonstrated how this can enable rapid determination of homo-oligomeric states of target MPs. Finally, using a heteromeric MP complex, we demonstrate how the database can guide the extraction of over-expressed MPs, and even multi-protein complexes, for rapid and efficient extraction into native nanodiscs for downstream structural/functional studies.

### High-throughput bulk membrane solubilization assay

We first sought to develop a rapid, scalable assay that can be performed in most laboratory setups to quantitatively determine the native-nanodisc forming capability of a target MAP against a specific cell type. Recent years have seen a rapid expansion of membrane-active co-polymers (herein referred to as MAPs, as a collective descriptor) proposed to form native nanodiscs. There are at least 30 commercially available, with many more proposed MAPs (Fig. 1a). Most often such membrane solubilization capabilities of MAPs are tested against simple synthetic liposomes, where a target MAP is incubated with synthetic liposomes and the solubilization is tracked by the dissolution of the parent liposome through light scattering approaches. But as shown recently, when tested against native cell membranes, such results are not always recapitulated^19^. Addressing this, our first goal was to develop a high-throughput assay that can be directly applied to native cell membranes to quantitatively report the native-nanodisc forming capability of a target MAP in a rapid and cell-type-specific manner. In our attempt to understand why polymers benchmarked using liposomes often fail to solubilize native membranes, we observed that scattering-based assays are the most widely used experimental approach to evaluate liposome solubilization capabilities of MAPs. These assays track the dissolution of the initial, larger parent liposome upon the addition of MAPs, with the assumption that any smaller-sized particles generated in the dissolution process are solubilized MAP discs. However, previous experimental and theoretical studies have also established that as MAPs commence the solubilization of the membrane, the resulting solution contains a mixture of both MAP-discs and smaller undissolved membrane vesicles ^24,25^. Therefore, these traditional scattering-based measurements that assume all smaller particles as discs inflate the actual % of the membrane scooped out as discs by the MAP. Consequently, when applied to the native membrane, MAPs would produce a significantly smaller % of nanodiscs. More importantly, in a native membrane context, the generated small vesicles would be recalcitrant to both standard affinity-enrichment or SEC approaches used to purify the target MP-containing nanodiscs, leading to large sample loss.

**Fig. 1:**
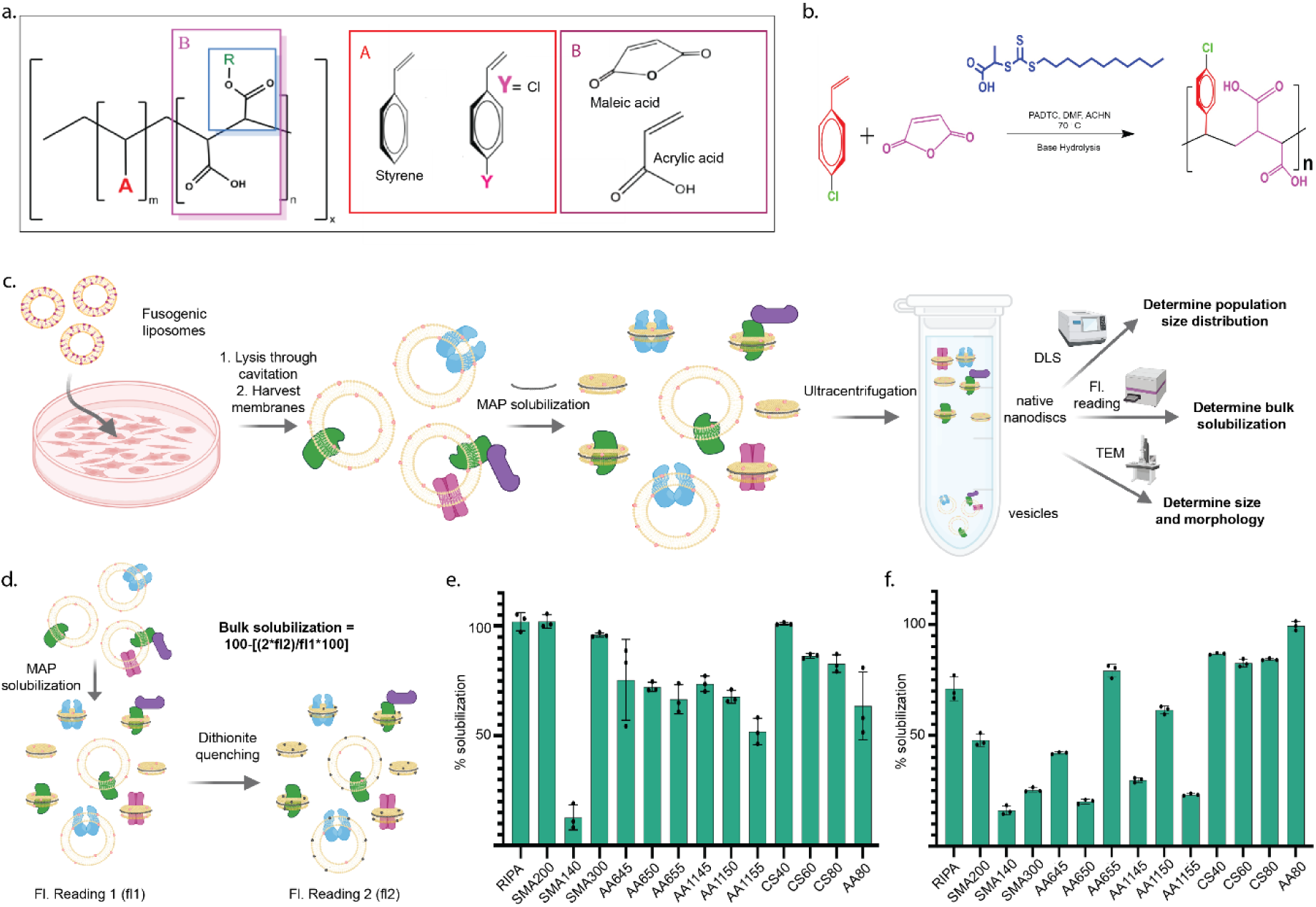
Generation of MAP library and development of high-throughput bulk solubilization assay. a, Representative chemical structures of commercially available and in-house synthesized membrane active polymers (MAPs). The diversity of MAP structures is manipulated by varying hydrophobic and hydrophilic moieties, their ratios, functionalization of R groups, and polydispersity. b, General synthesis scheme for RAFT-based polymerization of MAPs. c, Schematic of fluorescence bulk solubilization assay. Fusogenic liposomes are used to deliver fluorescent lipids to living cells labeling the membranes with a fluorophore. Membranes are then harvested, MAP solubilized, and subjected to a range of quality control experiments to determine bulk membrane solubilization through fluorescence measurements and size distribution and homogeneity through dynamic light scattering and negative stain EM d, Schematic of bulk solubilization quantitation. Membranes labeled with fluorescent lipids are solubilized with MAPs and a fluorescence reading (fl1) is taken before subsequent quenching with dithionite. Post-quenching, a second fluorescence reading (fl2) is taken. The precise nanodisc extraction efficiency is calculated using Equation 1. e, Bulk solubilization screen of polymer library against Hek293 cells and f, HeLa cells, where before solubilization fluorescent lipids were delivered to the cell through fusogenic liposomes. Subsequently, the cells were solubilized using respective MAPS and the % solubilizations were calculated using the procedure detailed in the panels c and d. Here 100% solubilization would indicate that the entire membrane was extracted into MAP discs, as 100% corresponds to the total starting fluorescence of the membrane. The error bars indicate the standard deviation across three solubilization replicates.

To overcome this, and benchmark our MAP library (Extended Data Fig. 2) in a biologically meaningful way, we developed a dithionite-based fluorescent quenching assay that can distinguish between MAP-solubilized native nanodiscs and un-solubilized small membrane vesicles ^26,27^, reporting the true membrane solubilization capability of a MAP in a quantitative manner. This is based on the principle that while dithionite can quench fluorescent lipids present on both bilayer leaflets of flat MAP discs, it only quenches 50% of vesicular fluorescent lipids located in the outer leaflet of the membrane. To calculate the percent solubilization into nanodiscs, membranes labeled with fluorescent lipids are solubilized with MAPs and a fluorescence reading (fl1) is taken before subsequent quenching with dithionite. Post-quenching, a second fluorescence reading (fl2) is taken. This value is multiplied by 2 to represent the solution fluorescence coming from vesicles. We then calculate the percentage solubilization into nanodiscs using Equation 1 which enables precise determination of the extent of membrane solubilization.

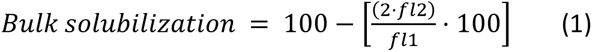

We first validated our approach using fluorescent liposomes as a model system (Extended Data Fig. 3). As expected, we observed that even while solubilizing liposomes, a significant portion of the membrane remains in undissolved small vesicle form that does not get quenched by dithionite, requiring at least 200,000g ultracentrifugation to be completely separated from discs.

We next modified this assay for direct application to mammalian cells (HEK293 and HeLa). Here, we leverage fusogenic liposome technology to deliver fluorescent lipids directly to target mammalian cell membranes^28^. Subsequently, the initial fluorescence of the cell membrane is recorded, before subjecting it to a comprehensive MAP solubilization screen where we systematically vary the polymer solubilization conditions. As with the fluorescent liposomes, the fluorescence of the post-solubilization membrane (after dithionite quenching) is compared with the initial membrane to yield the solubilization efficiency (Fig. 1c,d). Figure 1e-f shows the application of this on two different cell types using nine commercially available polymers, as well as five homemade MAPs. The homemade MAPs were synthesized in a chain-length controlled manner using established RAFT chemistry^29^ (see Online Methods for details). RIPA was used as a control. The presence of MAPdiscs in the solubilized sample was further confirmed by negative stain EM imaging (Extended Data Fig. 3). Figures 1e-f quantitatively show the bulk membrane solubilization % of each of the MAPs against each of the cell types. This provides a rapid and quantitative assay using which any existing or newly synthesized polymer can be assessed against a target cell line to quantitatively evaluate its bulk membrane solubilization capability, setting up a benchmark in both polymer and cell type-specific manner. For example, here we can see that both AASTY1155 and SMA140 perform poorly in bulk solubilization of the HEK293 cells. Consequently, they are not considered in subsequent experiments. For future work, we expect this fluorescence quenching-based native membrane bulk solubilization assay to be a community standard, replacing the typically used liposome scattering assay and eliminating any discrepancy in extraction efficiency.

### Membrane proteome-wide quantitative extraction database

The fluorescence-based screening method mentioned above, although useful for assessing the bulk solubilization properties of polymers in a cell-specific manner, lacks molecular resolution. Different polymers extract different amounts of a target protein into native nanodiscs. The dithionite assay cannot capture this variability at an individual protein level and hence, cannot indicate the MAP that is most efficient in extracting a target protein. Addressing this, we next developed a comprehensive membrane proteome-wide extraction database, which provides the most efficient extraction conditions for the MPs present in a specific cell type. Applying this to HEK293 cells we have developed a quantitative database of that captures variable extraction efficiency of 2065 unique MPs across 11 different MAP conditions. At an individual protein level, this indicates what would be the best MAP for efficient extraction of a target MP into native nanodiscs.

To accomplish this, we integrated MAP extractions with Label-Free Quantitative (LFQ) proteomics. We designed a workflow for membrane proteomics from MAPS modifying previous protocols of detergent-based proteomics (Figure 2a)^8,30,31^. The harvested membranes from the target cell type were subjected to a MAP solubilization screen with variations in the target polymers or solubilization conditions. After solubilization, the corresponding MAPdiscs were isolated via ultracentrifugation at 200,000 x g. To ensure sample quality, a small portion of these isolated MAPdiscs were subjected to Dynamic Light Scattering (DLS), negative stain Electron Microscopy (EM), and the fluorescence-based bulk solubilization screen (Extended Data Fig. 3). The remaining samples underwent LFQ-based quantitative proteomics analysis (Fig. 2a). This comprehensive experiment yielded relative abundance profiles for all the MPs detected in the cell type, under various extraction conditions. Compilation of this relative abundance for each MP, in each condition tested, yields a membrane proteome-wide extraction library capable of providing the most optimal extraction conditions for any of the MPs detected. We applied this to HEK293 cells, and based on our bulk membrane solubilization data, we identified the top 11 polymers exhibiting over 50% bulk solubilization efficiency for further investigation.

**Fig. 2:**
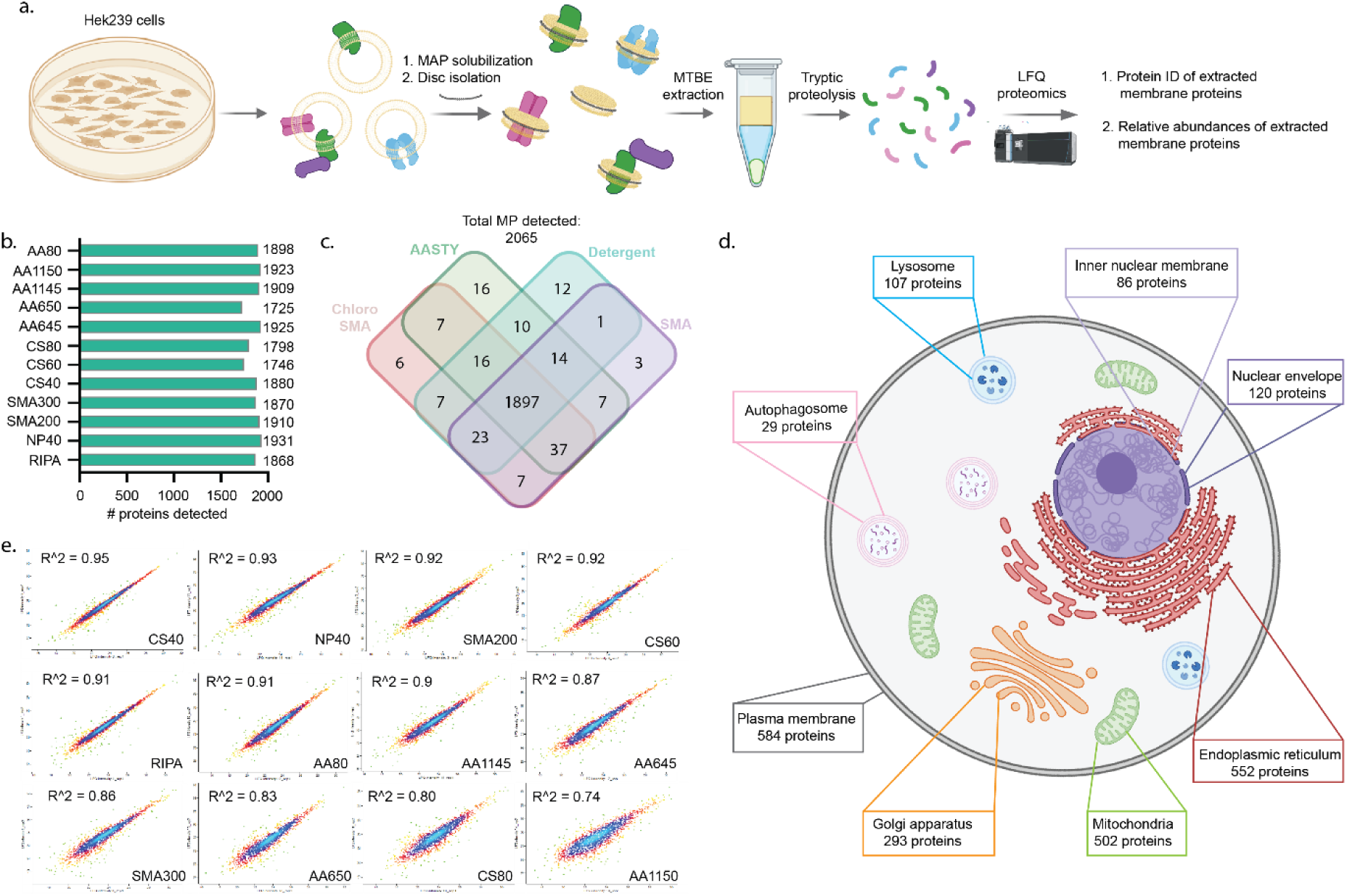
Integrated extraction workflow and proteome-wide qualitative assessment of MAP extraction. a, Schematic of experimental workflow for MAP extraction and proteomics analysis. Cellular membranes are solubilized with MAP library. From the MAPdiscs isolated through ultracentrifugation, proteins are precipitated using MTBE extraction and subsequently reduced, alkylated, and digested overnight with trypsin. The tryptic peptides are subjected to label free quantitative proteomics through LC-MS/MS. b, Number of MPs detected in each extraction condition providing an overview of efficacy of MAPs in extracting MPs. c, Overlap of protein coverage from different extraction conditions. The polymers are divided into three broad classes (SMA: SMA200, SMA300; AASTY: AASTY645, AASTY650, AASTY1145, AASTY1150, and ChloroSMA: ChloroSMA40, ChloroSMA60, Chloro80). As depicted, while there is an overall robust overlap between different conditions, 83 MPs are only detected under the MAP conditions. d, Membrane proteins are detected from all membrane-bounded cellular organelles yielding robust organellar coverage. Membrane proteins detected in the proteomic screen were filtered through organellar databases obtained from Uniprot, and only proteins with specified organellar residence were annotated, and proteins with more than one organelle of residence were annotated for each organelle. Organellar analysis reveals that MAP solubilization is not limited to the surface proteome as proteins were robustly detected from all endogenous intracellular organelles. e, Variation in label free quantitation was measured for each protein across each biological replicate. The biological replicates were plotted against each other to capture reproducibility of solubilization. Here, an R^2 value closer to 1 indicates better reproducibility. As shown, most MAPs show a very high degree of reproducibility. Interestingly, AASTY1150, which showed poor bulk solubilization, also showed poor reproducibility.

Our analysis (see Methods for details) yielded IDs of more than 4000 proteins, which includes both MP (integral and peripheral MPs), as well as soluble proteins associated with membranes through protein-protein interactions with other MPs (such as TRC40). From this, we then filtered the subset of the proteins designated as integral or membrane-associated proteins in Uniprot. As shown in Figure 2b, this yielded a comprehensive list of 2065 MPs that we can detect and further quantify their abundance across all the MAP extraction conditions. To date, this presents the largest extraction screen that provides the most in-depth quantitative view of spatial extraction of membrane proteomes. Our screening also included two of the most-used detergent conditions used for proteomics studies, RIPA and NP40, as controls. Surprisingly, in our optimized protocol (See Methods), most MAPs perform comparable to these detergents, in the number of MPs identified, in some cases out-numbering them. Using the LFQ intensities, we quantified the relative abundance of all 2065 MPs in each of the extraction conditions. As shown in Figure 2b, a significant number of MAPs outnumber RIPA in the number of MPs that are best solubilized (shows the highest LFQ intensity) in them. Overall, 1897 MPs could be detected and quantified in all conditions (Fig. 2c), with 83MPs that are only present in one or more MAPs and not in any detergent conditions. We further classified these proteins into their known organellar membrane of residence based on curated organellar databases from Uniprot. The data clearly demonstrates our ability to extract MPs from not just PM, but all endogenous organellar membranes, including ER, mitochondrial inner and outer membrane, nuclear inner and outer membrane, lysosome, Golgi, as well as transient organelles such as autophagosomes and endosomes (Fig. 2d). Each of the MAP solubilizations were replicated 8 times (2 Biological replicates X 4 technical replicates). Using this we further sought to understand the reproducibility of each polymer’s extraction capacity by measuring the R^2^ variation in the LFQ value for each MP across biological replicates (Fig. 2e). As shown, most polymers were found to be very reproducible in their total solubilization efficiency. Notably, AASTY1150, which showed poor bulk solubilization, also showed poor LFQ reproducibility.

Next, using this LFQ intensity, we build a database that reports the extraction efficiency of each of the 2065 MPs across all the tested MAP extraction conditions. For each target MP, the condition providing maximal extraction was treated as 100, all others were represented as relative to that. We have further built a web app through which this database can be quickly searched to obtain the most optimized conditions to extract a target MP into a native nanodisc (Other Supplemental Information, www. https://polymerscreen.yale.edu/). Figure 3a shows an output from the database for an example membrane protein, EGFR. Here, the gene-name search directly yields the relative extraction efficiency in all the MAPs within the database. This provides a quantitative guide that can directly yield the most optimized condition to extract a target MP from its physiological membrane into a native nanodisc that preserves the local membrane context. In the next section, we will provide some key examples of how this could be used to rapidly extract a target MP from its native membrane and perform downstream bioanalytical studies. One intriguing conclusion that stems from Figures 3c-e is MAPs do have preferences in extracting MPs. Such preferences are likely to arise from the differences in the chemistry of different MAPs that lead to differential extraction efficiency towards both individual proteins as well as the membrane environment where they reside. This is highlighted in the unbiased clustering analysis of all 2065 MPs (Fig. 3f). Strikingly, polymers with similar chemistries tend to have similar extraction trends such as many of the AASTY series of polymers clustering together. To better understand the protein-specificity and the organellar membrane specificity, we subdivided all detected MPs by polymers that provided the best extraction (as shown in Fig. 3b). We then stratified these MPs by their number of TM helices, molecular weight, as well as organellar membrane. As depicted in both Figures 3c and 3d, there is a clear trend observed for both properties. Together Figures 3c-e shows that the specific chemistry of the MAPs imparts intrinsic molecular propensities that tune their efficiency towards specific proteins and organellar membranes. It is conceivable that parameters beyond what is analyzed here may also influence such gradation in efficiency. We hope our open access data presented here, as well as future similar proteome-scale studies on other cell types and newer polymers, can enable the community to perform an in-depth, large-scale meta-analysis to glean out those specific chemical properties that result in such observed gradation in efficiency. This would enable the development of next-generation MAPs that can be used to make precision extraction in an organelle-specific or protein class-specific manner.

**Fig. 3:**
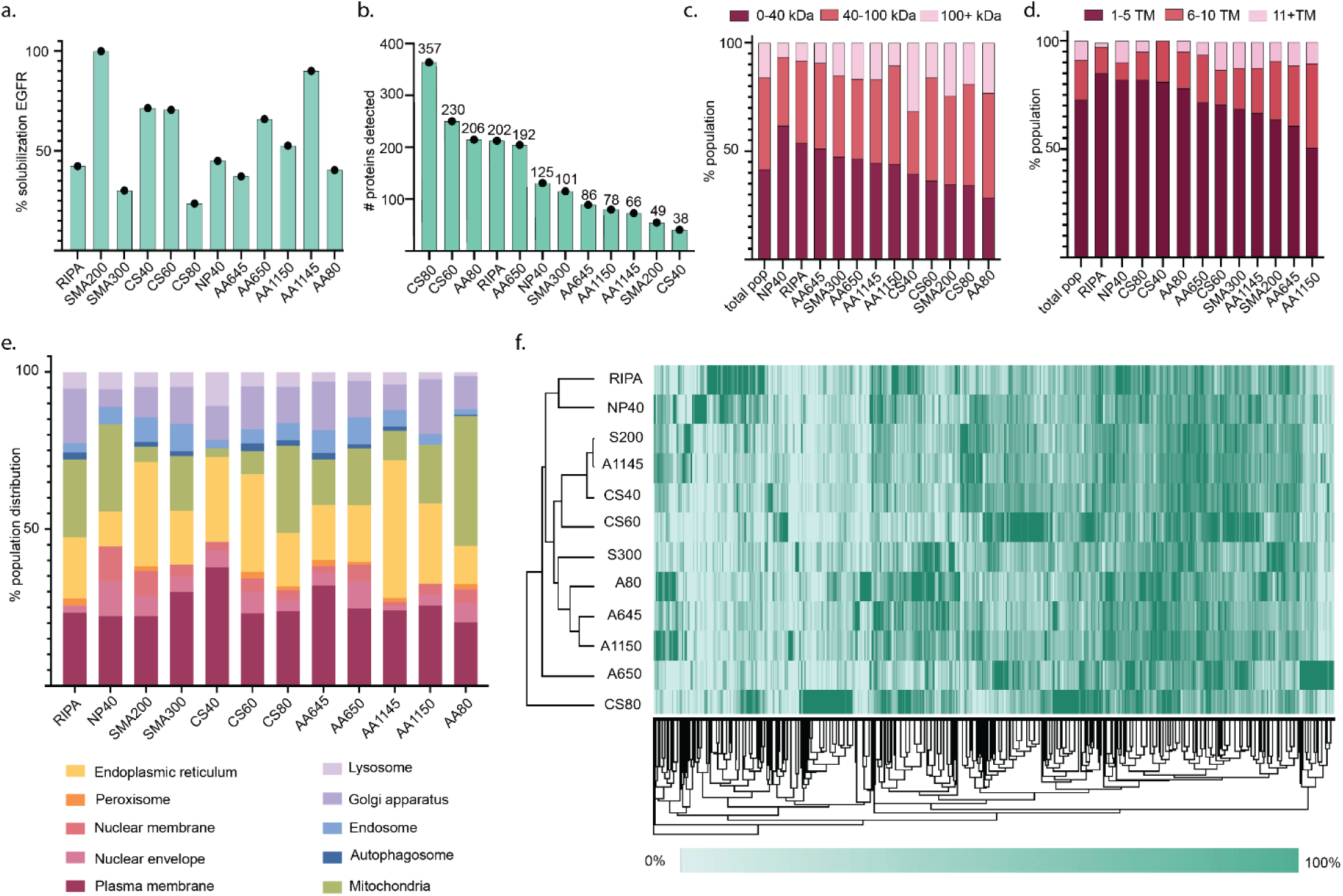
Quantitative analysis of MAP extraction efficiencies reveals solubilization preferences and polymer-specific trends. a, Solubilization plot from the open-source web application taking EGFR as a representative membrane protein. The Y axis shows the relative solubilization efficiency of EGFR under each MAPs used, as designated on X -axis. Based on the normalized solubilization efficiency, SMA200 is the optimal extraction condition for the protein. b, Number of unique MPs best solubilized under each MAP and detergent condition. For eg, for CS80, the number 357 denotes that CS80 is the best polymer for extraction for 357 unique MPs. As seen, three in-house synthesized polymers far outperformed commercial polymers in quantitative extraction efficiency. c, Stratification of the best solubilized proteins by molecular weight, reveals distinct population distributions. Detergents exhibit superior performance in extracting small molecular weight proteins (<40kDa), while MAPs demonstrate efficacy in solubilizing medium to large molecular weight proteins (>40kDa). d, Stratification of the best-solubilized proteins by number of transmembrane domains reveals distinct population distributions. Detergents preferentially solubilize proteins with small numbers of transmembrane helices (TMD, 1-5) while MAPs excel at extracting proteins with larger numbers of TMDs (>5). e, Detailed breakdown of organellar solubilization preferences across the MAP library, revealing distinct polymer-specific trends in extraction. f, Hierarchical clustering analysis provides grouping of MAPs by extraction efficiencies yielding a deeper understanding of polymer-specific solubilization trends.

### Validation and application of membrane protein spatial extraction database

#### A. Direct high-throughput extraction and purification of target individual MPs from organellar membranes

We next validated the general and broadscale applicability of our database and demonstrated how this can facilitate rapid extraction and purification of target MPs into native nanodiscs, directly from their physiological organellar membranes. For this, we selected a distinct set of MPs that are very diverse in their structure, functionality, as well as resident of different organellar membranes. This includes plasma membrane, Mitochondria, Golgi, ER, and lysosome. Concurrently, we established five distinct cells, expressing one of these organellar marker proteins at a level that ensures its retention within the target organellar membrane. Each of these proteins were tagged with eGFP to visualize and validate organellar localization through microscopy (Extended Data Fig. 4). Our MAP-extraction database provided the most efficient MAPs in extracting each of these proteins (Fig. 4a). Following this, we subjected the membranes of the individual cells to solubilization using the respective best-performing MAPs for each of the target MPs. The % efficiency of extraction was further calculated using the GFP-fluorescence (See Methods). As shown, in each case, we achieved the direct extraction of the target MPs into native nanodiscs from their resident organellar membranes (Fig. 4b). Further, the % solubilization data for each of these proteins highlights the high efficiency of extraction for each of these proteins. Taking the example of KRas, we further show that these target MP-containing native nanodiscs can also be affinity-purified to a very high degree of purity (Extended Data Fig. 4). This demonstrates how using the database as a quantitative guide, we can rapidly extract and purify target MPs with high efficiency, directly from their native membrane, in the form of a native nanodisc that preserves their physiological organellar membrane context.

**Fig. 4:**
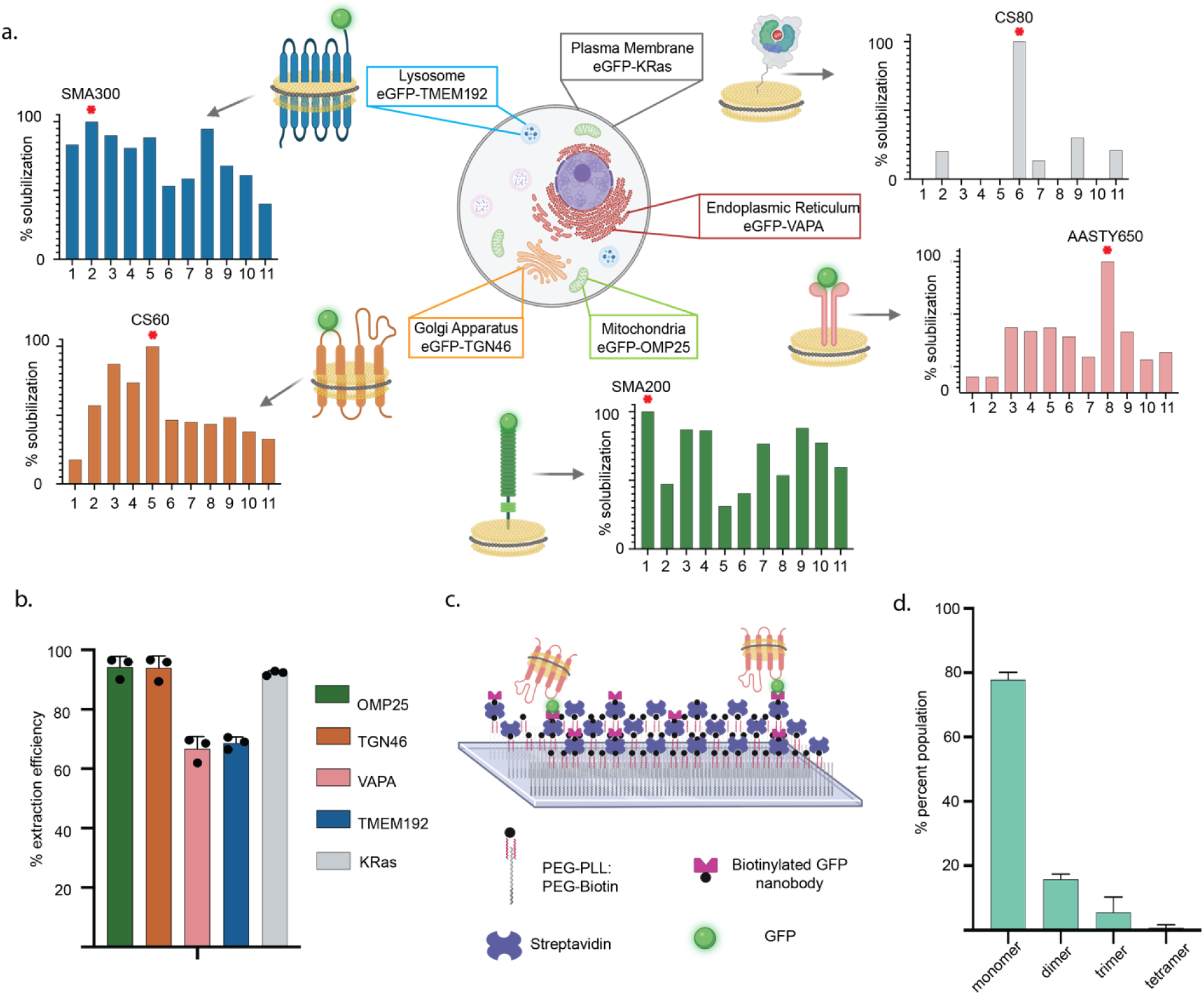
Database-driven extraction of intra-organellar membrane proteins and determination of oligomeric state through NativeNanoBleach analysis. a, Database provided optimal extraction conditions for 5 organellar MPs used as a benchmark. The “red-star” indicates the specific polymer that yields the best extraction. The corresponding best performing MAPs (indicated with red star) are chosen for extracting the target MPs b, Experimental extraction efficiency of each of the five GFP-tagged MPs extracted under the optimal MAPs, directly from their native organellar membranes to preserve native membrane environment (OMP25: SMA200, TGN46: CS60, VAPA: AASTY650, TMEM192: AASTY650, KRas: CS80). The error bars indicate the standard deviation across three solubilization replicates. c, Schematic of the NativeNanoBleach workflow. Glass slides are coasted with a PEG:PLL PEG:biotin conjugate. The surface is sequentially coated with streptavidin and a biotinylated GFP nanobody which immobilizes the GFP-tagged disc on the surface. This permits rapid isolation and immobilization of GFP-tagged target MP-containing discs directly from the MAP-solubilized lysate membrane, enabling downstream TIRF microscopy and step-photobleaching analysis d. Oligomeric distribution for GFP-TGN46 derived through nativeNanoBleach. The error bars indicated the standard deviation across three biological replicates for NNB analysis. The protein was extracted using CS60. Polymers from the database are 1: SMA200, 2: SMA300, 3: ChloroSMA20, 4: ChloroSMA40, 5: ChloroSMA60, 6: ChloroSMA80, 7: AASTY645, 8: AASTY650, 9: AASTY1150, 10: AASTY1145, 11: AASTY80

#### B. Determining endogenous oligomeric states

We next demonstrated an example of how the capability to rapidly isolate target MP containing native nanodiscs can be interfaced with orthogonal approaches to glean critical molecular information on membrane proteins. To do this, we couple our database with single-molecule TIRF microscopy to determine the homo-oligomeric states of MPs directly from their endogenous cellular membrane. To do this, we applied our recently developed NativeNanobleach approach^23^. Here, TIRF-based single-molecule GFP step-photobleaching of GFP-tagged target MPs in native nanodiscs yields their oligomeric states in the native membrane at single-molecule sensitivity. Further, it eliminates the need for prior purification of the target protein, as the glass slides are activated with GFP-nanobody which can capture the MAPdiscs containing GFP-tagged MPs (See Methods). This allows seamless integration with our high throughput database for rapid determination of endogenous homo-oligomeric states directly from the target physiological membrane. We demonstrate this using TGN46, a Golgi resident protein. As shown in Figure 4b, guided by the MAP database, we can directly extract GFP-TGN46 with 96% efficiency using ChloroSMA-60. We subsequently subjected these GFP-TGN46 MAP-discs to NativeNanoBleach. The MAP (ChloroSMA60 here) extracted total membrane was passed through glass slides conjugated with GFP-nanobody to capture the TGN46-MAPdiscs (Fig. 4c). Single-molecule step-photobleaching shows the presence of a distribution of the oligomeric state of TGN-46, ranging from monomer to tetramer (Fig. 4d). Indeed, recent in vitro work has indicated homo-oligomerization of TGN46 and its possible functional role in regulating the sorting of secretory proteins through the trans-Golgi network^32^. The data presented here offers an experimental avenue to study the link between in vivo oligomeric state and the functional role of TGN46; a work we are actively pursuing. But from a methodological perspective, this establishes an experimental pipeline, where using the MAP extraction database as a quantitative guide for optimal polymer selection, we can rapidly extract and determine endogenous oligomeric states of MPs directly from their native cell membranes.

#### C. Extraction and purification of target overexpressed MPs complexes

Finally, we demonstrate the application of our database in making informed decisions for the targeted extraction and purification of overexpressed multi-protein complexes. For this, we constructed a solubilization index, which represents a summed extraction efficiency of the target proteins in each tested polymer condition. To calculate the solubilization index, we sum the scaled LFQ values (x) representing the solubilization efficiency for each protein in the complex or pathway of interest. We then divide by the total number of proteins (n) effectively generating a solubilization average reflecting the total solubilization efficiency for the proteins of interest. The solubilization index is represented by Equation 2.

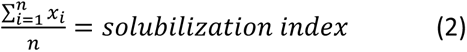

To validate the applicability of the approach, we focused on the VAMP2 and synaptophysin (Syp) complex system. VAMP2 is a single-pass transmembrane protein that binds to plasma membrane resident t-SNAREs effectively docking and fusing synaptic vesicles (SV) to the plasma membrane and regulating neurotransmitter release. Among the proteins present in SV, Syp, a multipass MP, is a major component whose exact function has remained elusive. Previous studies have suggested its complex formation with VAMP2, with low-resolution electron microscopy indicating potential higher-order multimeric structures^33–35^. However, challenges in the purification of both Syp, as well as VAMP-Syp complexes, have hindered direct observation and consequently, the determination of molecular organization and atomic structures for the possible Syp-VAMP2 complexes. The independent structure of Syp itself has also remained elusive. Using our MAP library, we sought to capture and purify the Syp-VAMP2 complex.

Based on the extraction efficiency data of the individual proteins in the database, we constructed a solubilization index plot that indicates that all ChloroSMA classes of polymers exhibit over 50% extraction efficiency, with ChloroSMA40 being the best-performing polymer (Fig. 5b). Notably, within the ChloroSMA-class we have observed ChloroSMA80 to produce larger-sized discs compared to other ChloroSMA classes of polymer (Fig. 5c). Taking into consideration the potential size of VAMP2-Syp macromolecular complex, we opted for the larger disc-sized ChloroSMA80 over the top-performing ChloroSMA40 of that class. We transiently transfected HEK293T cells with plasmids containing both Flag-tagged Syp and Strep-tagged VAMP2, subjected the membrane to extraction by ChloroSMA80, and subsequently purified the disc using a Flag antibody, followed by elution with the Flag-peptide. A Coomassie gel of the eluted sample confirmed the successful pull-down of the Syp-VAMP2 co-complex (Fig. 5d). The gel displayed two exclusive protein bands, with the top band corresponding to the mass of Syp and the lower band corresponding to the mass of VAMP2 (Fig. 5d). These identifications were further confirmed through both western blotting and GFP tagging of Syp (Extended Data Fig. 5). Since VAMP2 lacks a Flag tag, its presence in the pulldown disc sample can only be attributed to its co-complexation with Syp (Fig. 5a). We also confirmed that this co-purification was not a result of random protein association by overexpressing Syp alone, and subsequently extracting and purifying it using the ChloroSMA80 polymer and Flag resin (Extended Data Fig. 5). To our knowledge, this marks the first instance of a co-complex between different overexpressed integral-MPs being purified using any MAP. While this opens avenues to study the molecular and structural organization of the VAMP2-Syp complex directly from a membrane environment, it also provides a roadmap for extracting multiprotein complexes using our MAP library as a quantitative guide. Taking VAMP2 and Syp as a target system, we demonstrate how the database can be used to glean the most optimal condition for the extraction of specific protein complexes for downstream structural or functional studies.

**Fig. 5:**
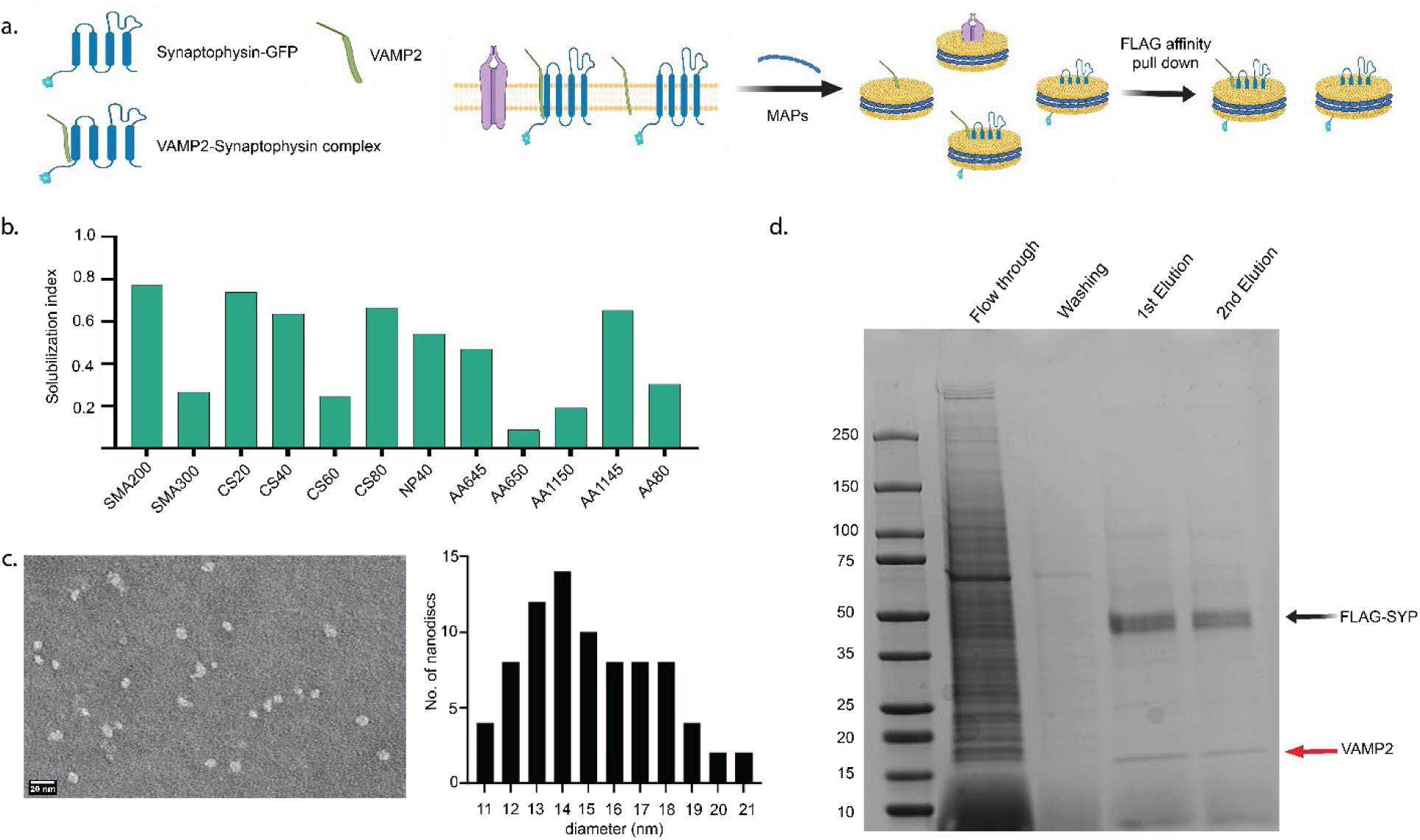
Native nanodisc extraction and purification of the synaptophysin-vamp2 co-complex. a, Schematic of FLAG-SYP-VAMP2 complex purification. FLAG-SYP and VAMP2 are coexpressed in EXP293T cells and solubilized using ChloroSMA80 as guided by the database solubilization index. MAPs containing FLAG-SYP are affinity purified using affinity pulldown. b, The calculated solubilization index for the SYP-VAMP2 complex. The CholorSMA series of polymers, boxed in pink, performed well as a group, with CS80 being chosen for downstream purification because of its propensity to form larger discs. c, Negative stain EM analysis of discs formed by CS80. Based on the population size distribution, CS80 forms discs of larger diameters than any other polymer in the CS series (See Extended Data Fig. 5). For the best chance of capturing the oligomeric complex of SYP-VAMP2, CS80 was chosen due to this propensity to form larger discs. d, Coomassie-stained gel shows the presence of 2 bands, one corresponding to the molecular weight of SYP and the second corresponding to the molecular weight of VAMP2. The presence of VAMP2, which does not have the Flag-tag, in the pulldown indicates successful co-complex purification.

## Discussion and Conclusions

In this study, we have developed a proteome-wide platform that provides a quantitative roadmap for the precise extraction and purification of target MPs, preserving their native spatial organization through the use of MAPs. Our publicly available database contains optimized extraction conditions for 2065 membrane proteins, offering a powerful tool for researchers in the field. Through the examination of five individual membrane proteins located in various cellular organelles (i.e. plasma membrane, endoplasmic reticulum, Golgi apparatus, lysosome, and mitochondria), as well as a multi-membrane protein complex, we validated how our database facilitates the rapid extraction and purification of target proteins directly from their physiological membrane environment. While this work has primarily focused on HEK cells for their widespread use, an obvious future goal is to extend these methodologies to other cell types, particularly those hosting different endogenous MPs. Efforts are underway to incorporate bespoke cell lines, such as neuronal or immune cell-lines, broadening the spectrum of proteins beyond those naturally occurring in HEK cells. Additionally, we are actively pursuing the expansion of the polymer library and the exploration of novel chemical conditions for extraction, such as varying pH and dielectric constants, present exciting opportunities to further enhance our spatial extraction capabilities. As and when these new conditions are tested, they can easily be integrated into the existing library, expanding its depth and breadth of coverage.

By integrating these approaches with orthogonal techniques, we anticipate a transformative impact on our understanding of membrane biology with unprecedented nanometer-scale resolution. This transcends the initially assumed roles of these polymers as mere vessels to do structural studies of over-expressed MPs. Our collaborative efforts have already demonstrated the potential of MAPs in elucidating the functional states of endogenous MPs and their influence on downstream cellular signaling pathways^23,36^. Furthermore, our development of native mass spectrometry platforms enables the direct detection of membrane protein-lipid assemblies from the membrane itself^37,38^. The integration of these orthogonal technologies with MAP-based extraction platforms facilitating the rapid extraction of target proteins and their native interactome holds immense potential across all facets of membrane biology. In this study, we have exemplified this potential by coupling MAP-based extraction with Native-nanoBleach, showcasing the platform’s ability to swiftly reveal endogenous homo-oligomeric states of MPs without the need for large-scale protein expression and purification. This serves as a compelling example of how complementary experimental approaches, when combined with database-guided extraction, can yield profound molecular insights into the structural and functional intricacies of MP-assemblies, directly from their native environment.

In addition to offering comprehensive extraction guidelines, we envision that the extensive proteome-scale data presented in this study holds the potential to inform the development of tailored next-generation MAPs. This notion is exemplified in Figure 3 c-f, which highlights the specificity of existing MAPs towards both organelles and specific proteins. Currently, the precise factors influencing why a particular MAP may perform more effectively for a specific protein, compared to others in the same cell, remains unclear. Plausibly, this preference may be rooted in the chemical affinity between a MAP and the target MP, the affinity between the MAP and the lipid domain housing the target protein, or potentially a combination of both. However, such nuanced insights are challenging to extract from small-scale studies focused on individual proteins with a limited selection of polymers. We believe that large-scale proteome-wide analyses against diverse MAP libraries can provide the reservoir of data necessary to elucidate these specific chemical interactions. Understanding these specificities holds tremendous promise in shaping the development of the next generation of MAPs tailored for organelle or protein-class-specific extractions. We are optimistic that our openly accessible data, beyond its role as a quantitative extraction guide, will catalyze the broader scientific community to advance in this pivotal direction.

## Author contributions

CB and KG conceived the study and designed the research. CB, with the help of MK, WL, and YL performed all the proteomics experiments. CB, with the help of RM and ES, established the online database. CB performed organellar extraction experiments. SG, with the help of TA, synthesized all the MAPs and established the bulk solubilization assay. SG, with the help of AP, SK, and JC, performed the experiments on the VAMP-Syp2 complex. JER provided guidance and insight on all experiments about VAMP2-Syp. GW performed all experiments and data analysis about Ras and NativeNanoBleach. MB provided guidance and insights on all experiments about Ras and NativeNanoBleach. KG and CB wrote the manuscript with contributions from all other authors.

## Acknowledgements

The VAPA, TMEM192, and TGN46 constructs were gifts from Dr. Pietro De Camilli, Dr. Shawn Ferguson, and Dr. Chris Burd and Dr. Julia von Blume respectively. We thank them for their generosity. This work in part supported by the National Institutes of Health grants RM1GM149406 and YALE POINTS award to KG and YL, R01GM141192 to KG, and R35GM147095 to MB. C.B. acknowledges support from the NSF GRFP fellowship (grant no. DGE-2139841) and the P.E.O. Scholar Award. The study was also funded by the joint efforts of the Michael J. Fox Foundation for Parkinson’s Research and the Aligning Science Across Parkinson’s Initiative. The Michael J. Fox Foundation for Parkinson’s Research administers the grant ASAP-000580 on behalf of Aligning Science Across Parkinson’s and itself.

## Data availability statement

All proteomics data is available via the PRIDE data repository at PXD052184. All remaining data will be made available from the corresponding author upon reasonable request.

## Materials and Methods

### Chemicals and Reagents

All reagents were purchased from Sigma-Aldrich and used without further purification unless specified otherwise.

### RAFT-based synthesis of in-house polymers

#### A. Synthesis

Cl SMA 40 Briefly, Chloro-styrene (388.2 µl, 3.1 mM), maleic anhydride (299.1 mg, 3.1 mM), and 2-(dodecylthiocarbonothioylthio) propionic acid (26 mg, 0.076 mM) were combined in a 50 mL Schlenk round bottom flask and dissolved in dimethylformamide. 1,1-Azobis(cyclohexanecarbonitrile) (10 mg, 0.004 mM) was added and dissolved.

Cl SMA 60 Briefly, Chloro-styrene (582.3 µl, 4.56 mM), maleic anhydride (450.5 mg, 4.56 mM), and 2-(dodecylthiocarbonothioylthio) propionic acid (26 mg, 0.076 mM) were combined in a 50 mL Schlenk round bottom flask and dissolved in dimethylformamide. 1,1-Azobis(cyclohexanecarbonitrile) (10 mg, 0.004 mM) was added and dissolved.

Cl SMA 80 Briefly, Chloro-styrene (776.36 µl, 6.1 mM), maleic anhydride (598.1 mg, 6.1 mM), and 2-(dodecylthiocarbonothioylthio) propionic acid (26 mg, 0.076 mM) were combined in a 50 mL Schlenk round bottom flask and dissolved in dimethylformamide. 1,1-Azobis(cyclohexanecarbonitrile) (10 mg, 0.004 mM) was added and dissolved.

The flask was capped with a rubber stopper and bubbled with nitrogen for 15 min. The solution was heated to 90 °C for 16 h. Polymers were purified by precipitation into either isopropanol or water, followed by filtering the polymers and drying in vacuo. A brownish solid polymer material was collected, weighed, and characterized by 1H NMR measured on a Bruker 400 MHz system (Extended data 2 and 6) and subjected to RAFT end group removal.

#### B. End-Group Removal of Poly (4 Chloro styrene-alt-maleic anhydride)

The dried polymer (ca. 4 g) was dissolved in dimethylformamide and combined with 2.4 g of benzoyl peroxide (9.9 mM) in a 250 mL round bottom flask. The flask was sealed with a rubber stopper and bubbled with nitrogen for 15 min. The escape needle was left in the flask, and the flask was heated to 90 °C for 6 h. Upon completion, the polymer was precipitated twice into either isopropanol or water, followed by filtering the polymers and drying in vacuo. Now this material was subjected to hydrolysis to produce Poly (4 Chloro styrene-alt-maleic acid).

#### C. Hydrolysis to produce Poly (4 Chloro styrene-alt-maleic acid)

Conversion of anhydride to acid was performed by hydrolysis in NaOH. Here 800 mg of polymer was dissolved in 2 g of KOH, 20 mL of water in a 100 mL round bottom and reflux the mixture for 4 h. This initially dissolved the polymer and at the end, reaction mixture becomes clear. The hydrolyzed polymer was subjected to dialysis using 3.5 KDa cut off membrane for 36 h. After that the polymer was precipitated by the addition of water and HCl(aq). Centrifuging the precipitate was followed by decanting off the supernatant. Wash the precipitate using 0.1 N HCl water three times and dry the precipitate using nitrogen.

Weight average (Mw) and number average (Mn) molar mass and dispersity Mw/Mn of each homemade polymer were determined by using size exclusion chromatography (Extended Data Fig 2 and 6).

#### D. NMR

For the NMR experiment, 0.5 mg of polymer material (Cl-SMA and AASTY 80) was dissolved in 400 microliters of either DMSO-D6 or CDCl3 and measured 1H peaks by using BRUKER 400MHz instruments. The NMR peaks are analyzed by using MNOVA software.

#### E. Size Exclusion Chromatography

Number-average (Mn) and weight-average (Mw) molar mass and dispersity (Đ = Mw/Mn) of copolymers were obtained from size exclusion chromatography (SEC) carried out using an AKTA Pure FPLC (GE) instrument outfitted with a Superdex 75 column, Cytvia. One milligram of each polymer was dissolved in 500 µl of 50 mM Tris, 150 mM NaCl, and pH 8.1 buffer. A similar buffer composition was used as an eluent at 0.5 mL/min flow rate at a 4-degree temperature. Dextran standards were used to calibrate the SEC system. Analyte samples at 2 mg/mL were filtered through a 0.2 mm PVDF membrane before injection (50 μL). All the data was processed by using Origin 2022 software.

The detailed synthesis protocol was deposited to protocols.io (DOI: dx.doi.org/10.17504/protocols.io.ewov19noylr2/v1)

### Expression and purification of VAMP2-Syp complex into native nanodiscs

#### A. Mammalian expression constructs

VAMP2 and Synaptophysin (Syp) genes were cloned in a T2A polycistronic vector (RRID: Addgene_196991) and the exact sequence of proteins is as follows – Syp-1xFLAG-T2A-Strep-VAMP2. This construct was expressed in ExpiHEK-293 cell cultures (RRID: CVCL_0045) using ExpiFectamine as a transfection reagent (Thermo Fisher). Briefly, thawed cells were passaged three times before transfection and were grown for 48 hours before being spun down and rinsed in ice-cold PBS.

#### B. Expression and solubilization of membrane proteins in native nanodiscs

Syp-VAMP2 expressed cells were resuspended in lysis buffer (*Table S3*) supplemented with protease inhibitor cocktail tablets (Pierce, Thermo) and lysed using nitrogen cavitation (600 psi, 15 minutes). Debris and cell nuclei were removed by a soft spin (1500 rcf, 10 min, 4 °C) and clarified lysate was ultracentrifuged (200K rcf, 1 h, 4 °C) to isolate the membrane fraction. Membranes were solubilized using a 1.5% MAP (CS80) in membrane resuspension buffer (*Table S3*) at 4°C. Solubilized membranes were then subjected to another round of ultracentrifugation to pellet any undissolved membrane, isolating soluble native nanodiscs in the supernatant.

#### C. Native nanodisc purification

Native nanodiscs containing Syp-VAMP2 were purified using affinity chromatography (Anti-FLAG beads, Sigma) at 4°C. Samples were adjusted with FLAG-binding, washed extensively with wash buffer, and eluted with elution buffer containing 3X FLAG peptide (10 µg/ml, SIGMA) using gravity flow columns. Buffer compositions were the same as membrane resuspension buffer. Purified nanodisc samples were subjected to further analysis. The detailed protocol was deposited to protocols.io (DOI: dx.doi.org/10.17504/protocols.io.4r3l2qo24l1y/v1)

### Membrane Protein Extraction and Fluorescence-SEC Analysis

30 million of EXP293T cells expressing a GFP-tagged membrane protein (GFP-SYP-VAMP2) were resuspended in 3 ml of 1.5% polymer in TBS (50 mM Tris.HCl, pH 8 and 150 mM NaCl) supplemented with EDTA-Free Protease Inhibitor Cocktail tablets (Thermo Fischer). Next this whole suspension was incubated on a rolling table for 2 h. Large aggregates were removed from the suspension by ultracentrifugation at 200 000g for 45 min and 200 μL of the supernatant was loaded onto a Superose6 column (10/300 GL) pre-equilibrated with TBS buffer. Separation was performed at a flow rate of 0.5 mL/ min, and the eluent was detected by using a Shimadzu fluorometer with excitation at 488 nm, emission at 509 nm, and a recording time of 20 min. Eluent was collected for further analysis. The detailed protocol was deposited to protocols.io (DOI: dx.doi.org/10.17504/protocols.io.14egn676ml5d/v1).

### Transmission Electron Microscopy

The morphology and size of the nanodiscs were further investigated by negative stain transmission electron microscopic (TEM). Fluorescence-SEC eluent samples were prepared by diluting 1:10 and 1:100, in a corresponding buffer. Carbon-coated copper grids (200 mesh) were glow-discharged for 30 s. Sample (5 μL) was applied to the grid and after 60 s blotted off with Whiteman (ashless) filter paper and subsequently washed once, with uranyl formate (UF, 2% in H2O), and then finally stained with UF for 60s and blotting dry using Whiteman. Micrographs were taken on a JEOL JEM 1400PLUS electron microscope using an operating voltage of 80 kV. The average size of the nanodiscs was estimated from at least 30 or more well defined individual particles. The detailed protocol was deposited to protocols.io (DOI: dx.doi.org/10.17504/protocols.io.x54v92ydml3e/v1).

### Step centrifugation assay

HEK293 cells were grown to 90% confluency and harvested. Cells were resuspended in a cell lysis buffer (50 mM Tris pH 7.4, 150 mM NaCl) and subjected to nitrogen cavitation lysis at 750 PSI for 15 minutes. A “soft” spin was conducted at 4000 rpm for 10 minutes to pellet cell debris. The samples were then subjected to a series of sequential ultracentrifugation steps at the speeds of: 20,000 g, 100,000xg, 150,000xg, and 200,000xg, with 1 hour for each spin. After each spin, the sample was subjected to dynamic light scattering analysis to calculate population size distribution. The detailed protocol was deposited to protocols.io (DOI: dx.doi.org/10.17504/protocols.io.36wgqn75xgk5/v1).

### Bulk solubilization assay

HEK293 cells were grown to 90% confluency. Fusogenic liposomes were used to deliver fluorescent lipids to native membranes as described previously^28^. Briefly, lipids DOPE, DOTAP, and rhodamine-PE were mixed in a 1/1/0.1% ratio and dried under nitrogen. The lipids were resuspended in a 20mM HEPES buffer to a stock lipid concentration of 2.1 mg/mL. The solution was vortexed and sonicated for 20 minutes to form liposomes. The liposomes were then diluted 1:100 in cell culture media. Normal growth media was removed from the cells and replaced with growth media containing fusogenic liposomes (5mL volume for a 10cm plate). Plates were incubated at 37C for 20 minutes before media was removed and cells were harvested normally. Cell pellets were resuspended in cell lysis buffer (50 mM Tris pH 7.4, 150 mM NaCl) and subjected to nitrogen cavitation lysis at 750 PSI for 15 minutes. A “soft” spin was conducted at 4000 rpm for 10 minutes to pellet cell debris. Membranes were collected at 200,000xg for 1 hour, and the resulting pellet was directly resuspended into extraction buffer. The buffer conditions for each extraction condition are outlined in SI Table 3. Each sample was homogenized and rotated at 4C for 2 hours. A small aliquot of each sample was saved for fluorescence quantification, and a post-solubilization ultracentrifugation spin was conducted at 200,000g for 1 hour. A plate reader was used to take pre and post centrifugation fluorescent measurements, and solubilization efficiency was calculated accordingly.

### Proteomics experiments and development of the DATABASE

#### A. Sample preparation for LC-MS

Solubilized MAPs were MTBE extracted with minor modifications from previously described^39^. Briefly, 1 mL MTBE:MeOH (3:1) was added to 200uL MAP discs and vortexed for 45 minutes at 4C. Samples were sonicated for 15 minutes at 4C, and 650 uL of H_2_O:MeOH (3:1) with 1% KCl was added to the samples. They were vortexed, and centrifuged at 20,000xg for 5 minutes to precipitate protein. The supernatant was discarded, and the protein pellet was dried under vacuum. Protein was resuspended in 10M Urea and heated for 30 minutes. 10mM DTT was added to each sample and incubated at 56C with shaking for 1 hour. 20mM IAA was added to each sample and incubated at 22C with shaking for 1 hour in the dark. The urea concentration was diluted to 1.5M with 50 mM ammonium bicarbonate, and 1ug of trypsin was added to each sample and incubated overnight at 37C with shaking. Samples were desalted using SepPak18 cartridges, and a BCA assay was used to quantify peptides before drying under vacuum.

#### B. Liquid chromatography

Samples were resuspended in water + 0.1% formic acid and equally loaded at 500 ng for each run. Chromatography was conducted on a Vanquish Neo nLC with a home-packed C18 column (15cm x 75 uM ID) for separation of peptides by hydrophobicity. A 75-minute gradient was used with a buffer system of water/acetonitrile with 0.1% formic acid. The detailed protocol was deposited to protocols.io (DOI: dx.doi.org/10.17504/protocols.io.e6nvw1k97lmk/v2).

#### C. Mass spectrometry

Mass spectrometric acquisition was performed using a DDA method on the Orbitrap Eclipse platform (Thermo Fisher Scientific). MS1 scans were acquired using the Orbitrap detector at a resolution of 120000 and stored in profile mode. A mass range of 350-1400 m/z was used with an intensity threshold of 1.0e3. A cycle time of 1 second was used for ion isolation and MS2 fragmentation. Ions were isolated with a 1.4 m/z window and fragmented with CID energy at 30% activation. Product ions were detected at the ion trap set to rapid detection mode. Precursor ions were dynamically excluded for 20 seconds after one instance of detection. The detailed protocol was deposited to protocols.io (DOI: dx.doi.org/10.17504/protocols.io.e6nvw1k97lmk/v2). LC-MS data is available for download in the PRIDE Repository: PXD052184.

#### D. Raw proteomic data-processing

MaxQuant version 2.3.1.0 (RRID: SCR_014485) was used to ID peptides and perform label free quantitation^40^. Spectra were first searched against the SwissProt Human Proteome, downloaded from Uniprot (03/09/23). Modifications were set to Oxidation (M), Acetyl (Protein N-term), and Deamidation (N). Trypsin/P was used for digestion with a max number of missed cleavages set to 2. The minimum peptide length was set to 7. Match between runs was enabled. Label-free quantitation was enabled with the LFQ min ratio count set to 1, and FastLFQ was used. Peptide and protein FDRs were both set to 1%. The “protein groups” table was used for further analysis, and decoys and contaminants were filtered out.

#### E. Data analysis for building a solubilization database

Any proteins not detected in both biological replicates were filtered out. The protein groups sheet was then hand-curated to filter out any soluble proteins that were erroneously filtered into the membrane protein database. The protein groups sheet was then run through an in-house code that matched protein hits against organellar databases downloaded from Uniprot (03/09/23). Any protein not matching an organellar database was hand-curated to either be included in the correct organellar database or filtered out with the resulting sheet containing only membrane proteins. The resulting organellar membrane protein databases are available in Supplementary Information 2. This sheet, containing 2065 proteins, was used to generate the solubilization database and for all downstream analyses. It was run through an in-house code to average LFQ values across biological replicates, normalize LFQ values, and normalize each protein to the highest LFQ value across all polymer conditions. This means that the polymer with the highest LFQ value for a given protein was scaled to 100% and all other conditions were scaled accordingly. These values represent the solubilization database and were fed into the WebApp for searching.

#### F. Developing the WebApp database

Python version 3.10.5 (RRID:SCR_008394) scripts were written to filter proteomics data. To map identified proteins to respective organelles, proteomes from each organelle were downloaded and curated from UniProt identifiers. The proteomics data was matched to these organelle-specific databases and assigned to organelle(s) based on the gene name. To normalize the proteomics data, each gene ID and its label-free quantitation (LFQ) values were normalized so that the highest LFQ in any sample was set to 100% solubilization. This script was used to generate the normalized data used on the website.

Leveraging the Python code, which retrieves specific protein-related data from a proteomics database, we developed a web application designed to enhance user interaction and data visualization. Utilizing the Streamlit framework version 1.34.0 (RRID:SCR_024354), we integrated the direct embedding of the proteomics database and associated Python code into the application’s backend. This provided a robust foundation for the front end, where users can interact with the software intuitively. The application’s dynamic page structure responds to user inputs, dynamically adjusting content and functionalities based on the user’s preferences and actions. In addition, we crafted both a homepage and a general methods section, both aimed at providing users with clear instructions and insight into the application’s capabilities and underlying methodologies. The website code is available at https://github.com/eric-sun92/cell_lab.

### Database-driven organellar protein solubilization

Membranes from HeLa cells (RRID: CVCL_0030) expressing GFP-TMEM192, GFP-TGN46, GFP-OMP25, GFP-VAPA, and GFP-KRas were collected and solubilized with AASTY650, CS60, SMA200, AASTY650, and CS80 respectively. A small aliquot of each sample was reserved for quantification. Post-solubilization, samples were spun at 200,000xg at 4C for 1 hour to remove insoluble materials. GFP fluorescence readings were taken pre and post-solubilization and solubilization efficiencies were calculated accordingly. The detailed protocol was deposited to protocols.io (DOI: dx.doi.org/10.17504/protocols.io.36wgqn735gk5/v1).

### Native Nanobleach for determination of hierarchical organization of protein complex

This method was adapted from Walker, Brown et al.^23^ Briefly, flow chambers were created using functionalized glass slides. The chambers were prepared by cleaning glass slides, followed by treatment with poly-L-lysine (PLL) PEG and PEG-Biotin. Streptavidin and biotinylated GFP-nanobody were sequentially applied to the flow chambers. Finally, native nanodiscs were added and through the GFP-GFP nanobody interaction immobilized on the glass surface. TIRF microscopy was used for imaging with three-color acquisition to assess molecule density and confirm the lack of background signal. Photobleaching experiments were conducted using specific laser powers and exposure times. The acquired images were subjected to detailed analysis, which included single particle tracking using ImageJ software version 1.53c (RRID:SCR_003070) and custom MATLAB code version R2023a (RRID: SCR_001622). The resulting data, encompassing 1,000 to 1,500 individual particles for each biological replicate, were utilized to determine the population distribution of the oligomeric state. This analysis accounted for the GFP maturation efficiency and involved multiple biological replicates for each experimental condition, ensuring a comprehensive and robust assessment of molecular behavior. The detailed protocol was deposited to protocols.io (DOI: DOI: dx.doi.org/10.17504/protocols.io.ewov19nbklr2/v1.

## Extended Data Figures

**Extended Data Fig. 1:**
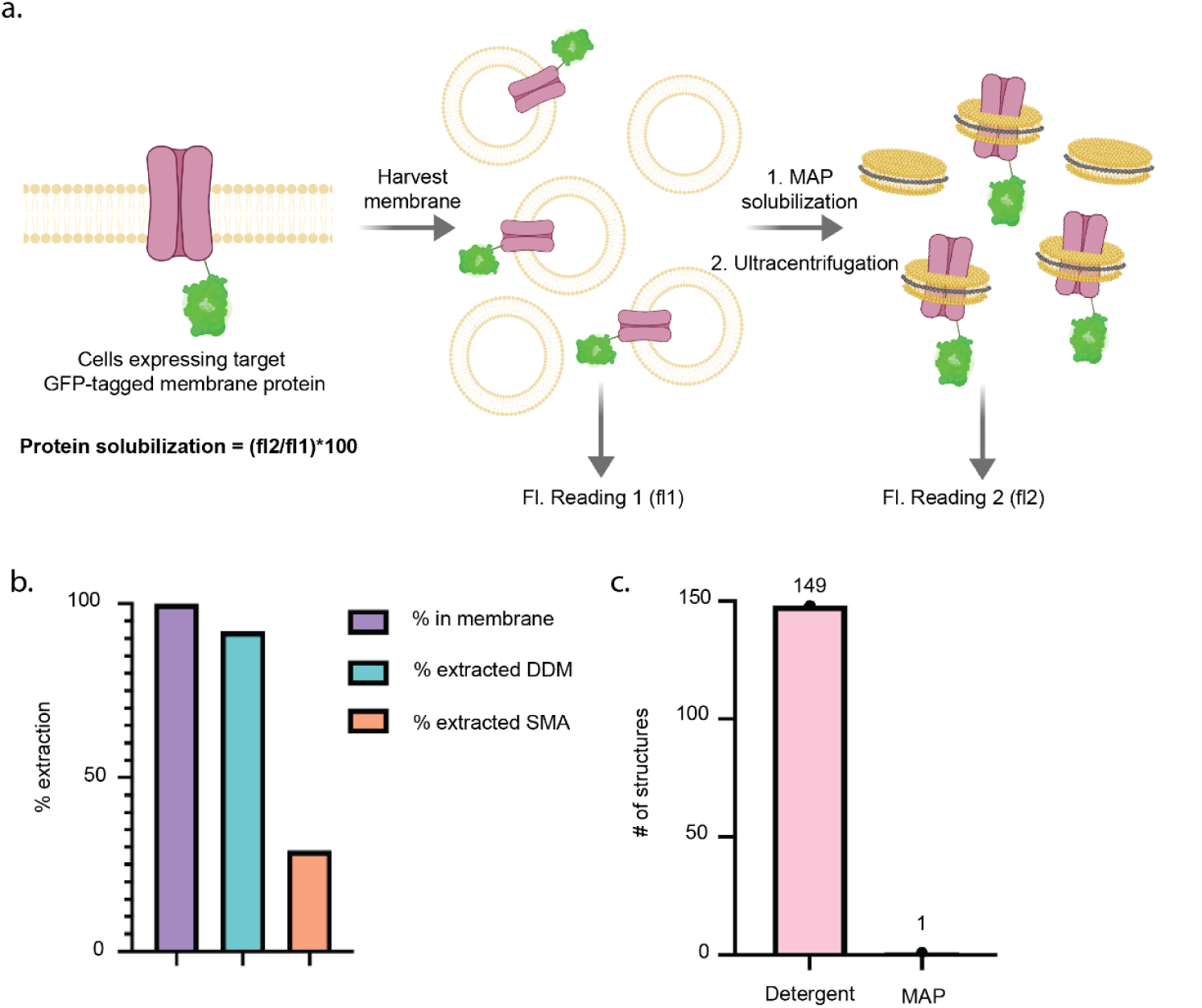
Comparative extraction efficiency of AqpZ and MAP extracted membrane protein structures: a, Schematic detailing GFP-based calculation for protein-specific solubilization efficiency. Membranes expressing a GFP-tagged target protein are harvested and an initial fluorescence measurement (fl1) is taken. Membranes are MAP solubilized and subjected to a 200,000g ultracentrifugation to remove insoluble material, and a second fluorescence measurement (fl2) is taken. Quantitative protein solubilization is calculated as the ratio of fl2:fl1. B, Extraction efficiency of model membrane protein Aquaporin Z into a traditional detergent system using widely cited polymer SMA200. Using standard conditions from the literature, SMA200 solubilizes poorly at 29% while standard detergent DDM solubilizes at 91%. B, Analysis of the last 150 deposited membrane protein structures to the PDB (9/7/23) reveals that only 1 structure was solved using polymer extraction. This highlights the current extremely limited utility of these native nanodisc-forming polymers which serve as a last resort for structural determination when other methods have failed.

**Extended Data Fig. 2:**
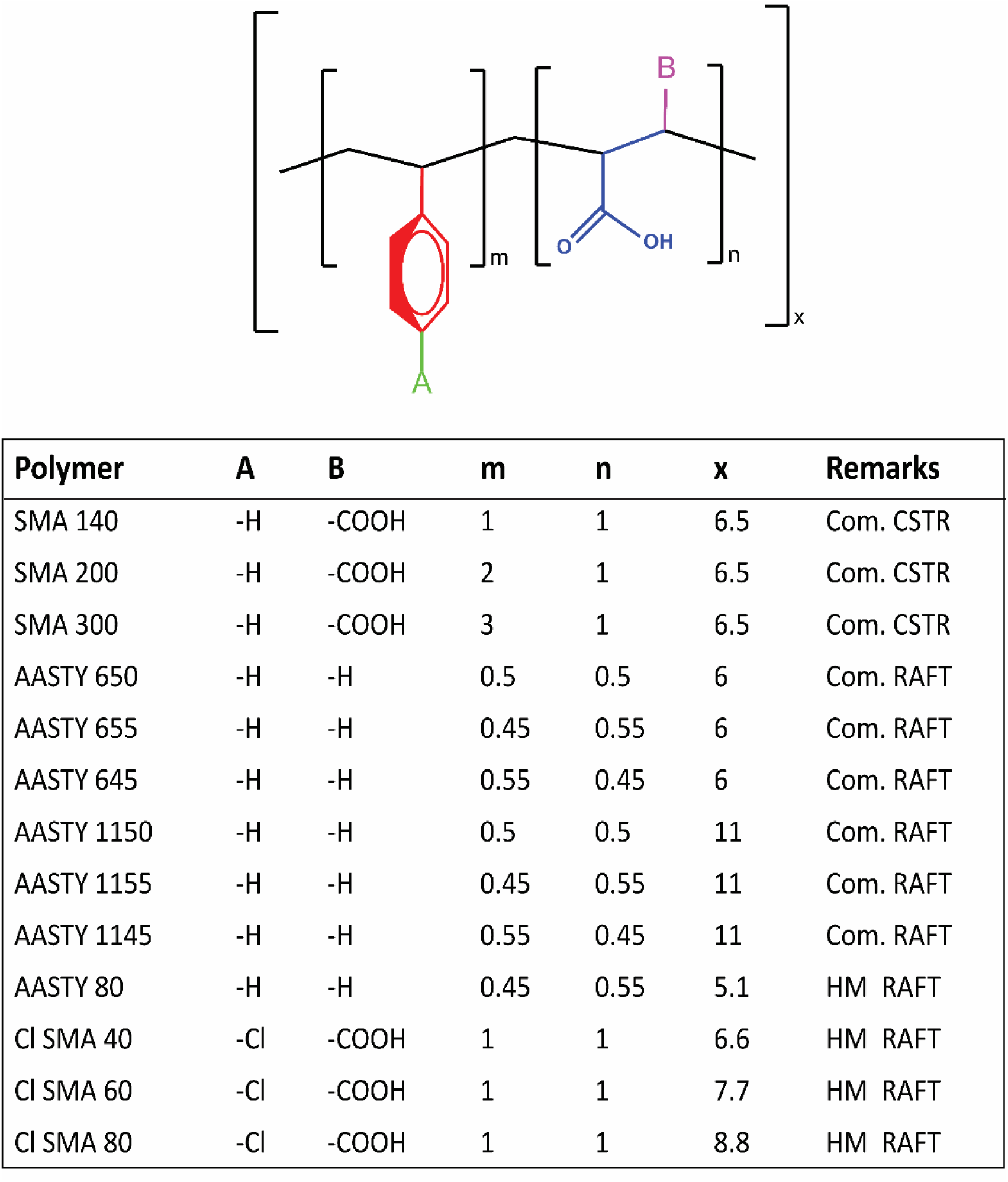
General Polymer structure. Table to show commercial (Com.) and homemade (HM) polymers. RAFT is noted for polymers that are synthesized using random addition fragmentation technology and CSTR is noted for polymers that are synthesized using continuous stirring tank reactor technology.

**Extended Data Fig. 3:**
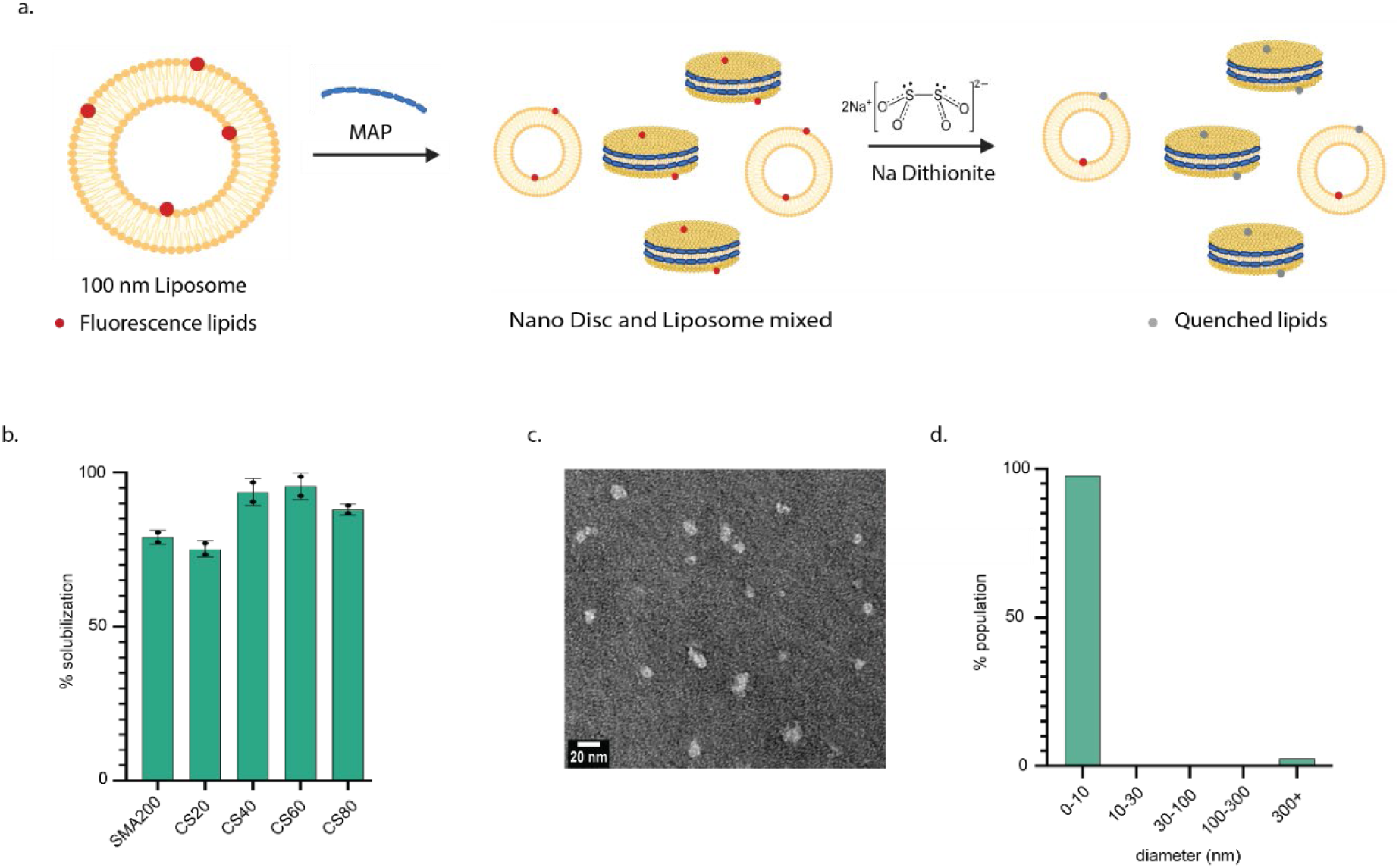
In vitro characterization of CS polymer series and quality control of samples through the proteomic pipeline. a, Bulk solubilization of fluorescent liposomes in vitro to characterize solubilization efficiency of in-house CS series of polymers. B, Representative negative stain EM image of discs in the preparation pipeline for proteomic analysis. C, Representative dynamic light scattering analysis of discs in the preparation pipeline for proteomic analysis.

**Extended Data Fig. 4:**
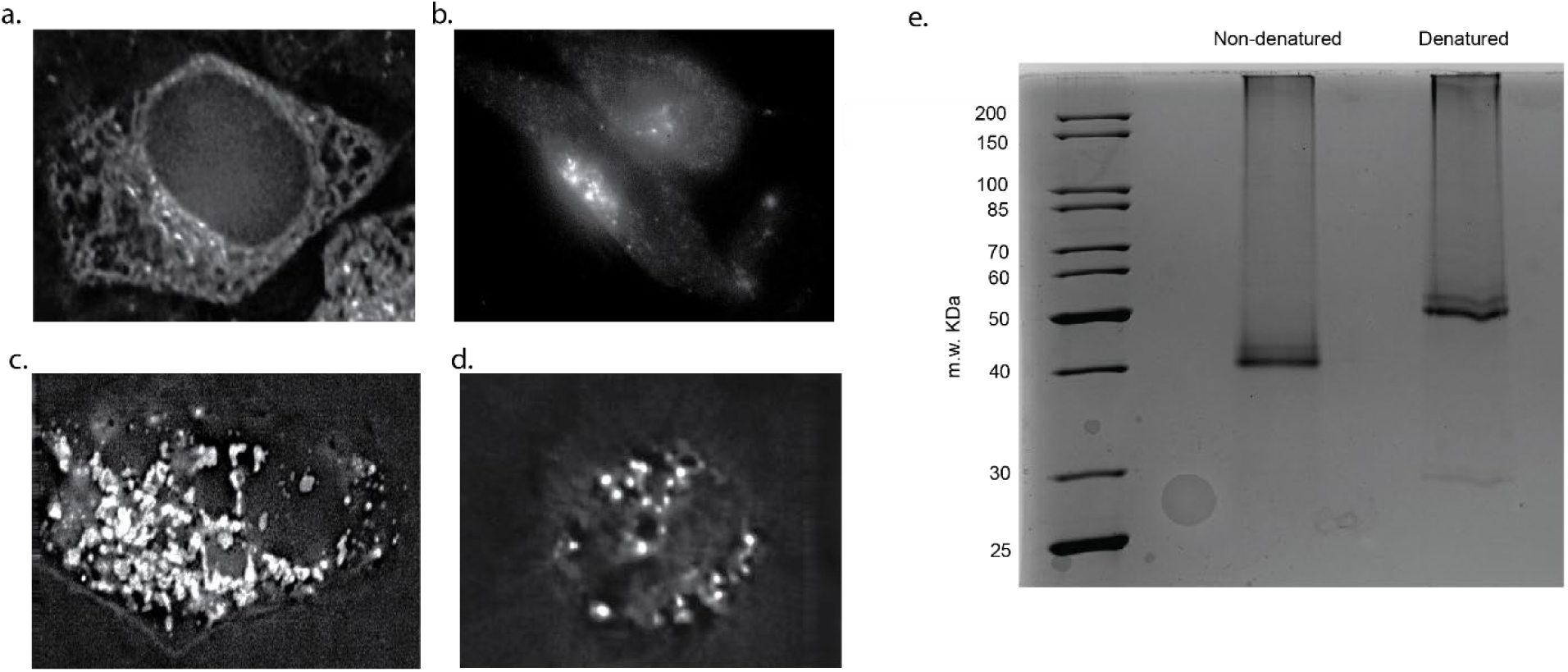
Confirmation of organellar localization and Kras purification. a-d, Deltavision images showing expression of organellar marker proteins is tuned to stay within their respective organelles. A, VAPA: endoplasmic reticulum b, TGN46: Golgi apparatus c, OMP25: mitochondria. This confirms a low level of expression for each of these organellar markers ensuring they remain localized within the target organelle. D, TMEM192: lysosome e, Coomassie stained gel of purified Kras in native nanodiscs. This gel shows a pure protein band corresponding to the molecular weight of peripherally associated membrane protein Kras showing a successful database-guided purification.

**Extended Data Fig. 5:**
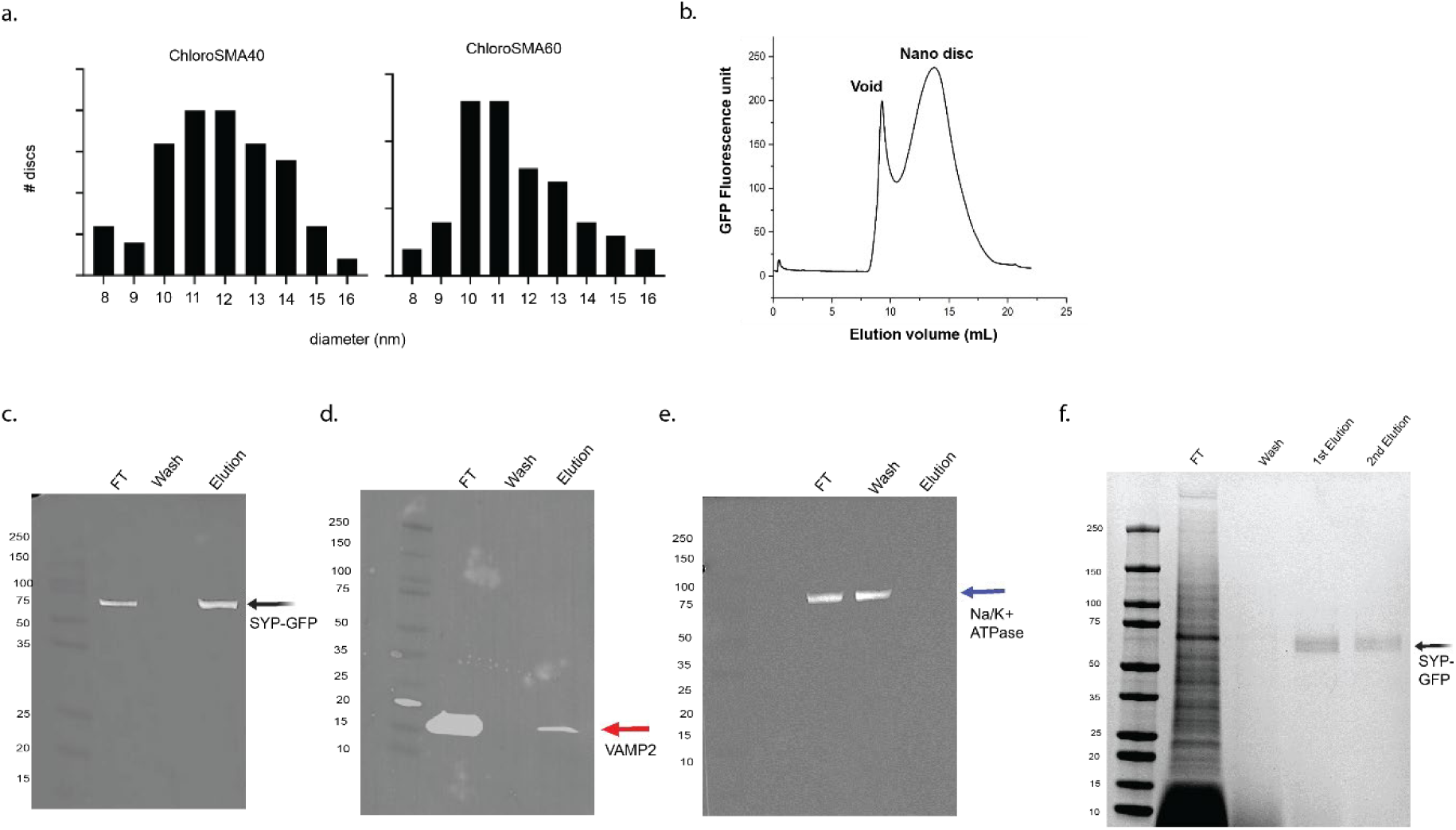
Characterization of purified SYP-VAMP2 co-complex. a, Population distribution of disc size across CS series. b, Fluorescence size exclusion chromatography of purified SYP-VAMP2 complex shows the purified product is in a stable nanodisc. c-e, Western blots confirm the identity and purity of SYP-VAMP2 complex. f, Coomassie gel from control experiment showing SYP purified in CS80.

**Extended Data Fig. 6:**
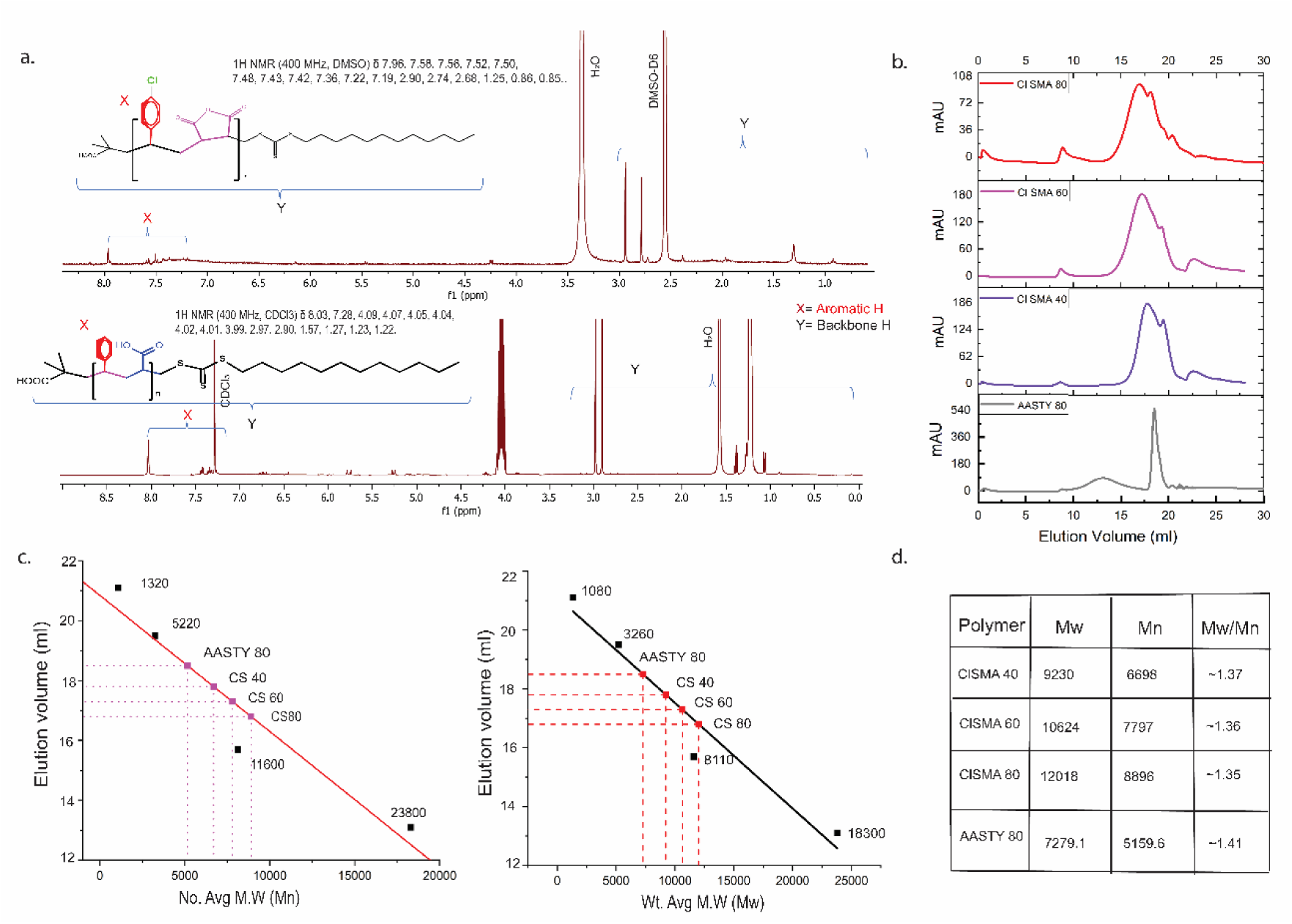
Characterization of Cl-SMA and AASTY 80. a, 1H NMR of Cl-SMAnh in DMSO-D6 and AASTY 80 in CDCl_3_. b, Size exclusion chromatography profile of homemade polymer AASTY 80, CS40, CS60, and CS80 ran through Superdex 75 column. c, Graph and d, Table stating the value of the weight average (Mw) and number average (Mn) molar mass and dispersity Mw/Mn of each homemade polymer. Dextran molecules of different molecular weight of were run as a standard to generate the calibration curve (represented as black dots in c).

## REFERENCES

1. Hakhverdyan, Z. et al. Dissecting the structural dynamics of the nuclear pore complex. Mol. Cell 81, 153–165.e7 (2021).

2. Pumroy, R. A., Fluck, E. C., Ahmed, T. & Moiseenkova-Bell, V. Y. Structural insights into the gating mechanisms of TRPV channels. Cell Calcium 87, 102168 (2020).

3. Kumar, A., Basak, S. & Chakrapani, S. Recombinant expression and purification of pentameric ligand-gated ion channels for Cryo-EM structural studies. Meth. Enzymol. 652, 81–103 (2021).

4. Levental, I. & Lyman, E. Regulation of membrane protein structure and function by their lipid nano-environment. Nat. Rev. Mol. Cell Biol. 24, 107–122 (2023).

5. Lundbaek, J. A., Collingwood, S. A., Ingólfsson, H. I., Kapoor, R. & Andersen, O. S. Lipid bilayer regulation of membrane protein function: gramicidin channels as molecular force probes. J. R. Soc. Interface 7, 373–395 (2010).

6. Reading, E. et al. The Effect of Detergent, Temperature, and Lipid on the Oligomeric State of MscL Constructs: Insights from Mass Spectrometry. Chem. Biol. 22, 593–603 (2015).

7. Urner, L. H. et al. Rationalizing the optimization of detergents for membrane protein purification. Chem. Eur. J 29, e202300159 (2023).

8. Nagaraj, N., Lu, A., Mann, M. & Wiśniewski, J. R. Detergent-based but gel-free method allows identification of several hundred membrane proteins in single LC-MS runs. J. Proteome Res. 7, 5028–5032 (2008).

9. Lee, S. C. et al. A method for detergent-free isolation of membrane proteins in their local lipid environment. Nat. Protoc. 11, 1149–1162 (2016).

10. Knowles, T. J. et al. Membrane proteins solubilized intact in lipid containing nanoparticles bounded by styrene maleic acid copolymer. J. Am. Chem. Soc. 131, 7484–7485 (2009).

11. Prakash, P. et al. Dynamics of Membrane-Bound G12V-KRAS from Simulations and Single-Molecule FRET in Native Nanodiscs. Biophys. J. 116, 179–183 (2019).

12. Routledge, S. J. et al. Ligand-induced conformational changes in a SMALP-encapsulated GPCR. Biochim. Biophys. Acta Biomembr. 1862, 183235 (2020).

13. Sander, C. L. et al. Nano-scale resolution of native retinal rod disk membranes reveals differences in lipid composition. J. Cell Biol. 220, (2021).

14. Dilworth, M. V., Findlay, H. E. & Booth, P. J. Detergent-free purification and reconstitution of functional human serotonin transporter (SERT) using diisobutylene maleic acid (DIBMA) copolymer. Biochim. Biophys. Acta Biomembr. 1863, 183602 (2021).

15. Mueller, S., et al. The bigger picture: global analysis of solubilization performance of classical detergents versus new synthetic polymers utilizing shotgun proteomics. *BioRxiv* (2023) doi:10.1101/2023.07.11.548597.

16. Sun, C. et al. Structure of the alternative complex III in a supercomplex with cytochrome oxidase. Nature 557, 123–126 (2018).

17. Qiu, W. et al. Structure and activity of lipid bilayer within a membrane-protein transporter. Proc Natl Acad Sci USA 115, 12985–12990 (2018).

18. Danielczak, B., et al. A Bioinspired Glycopolymer for Capturing Membrane Proteins in Native-Like Lipid-Bilayer Nanodiscs. BioRxiv (2021) doi:10.1101/2021.03.31.437849.

19. Smith, A. A. A. et al. Lipid nanodiscs via ordered copolymers. Chem (2020) doi:10.1016/j.chempr.2020.08.004.

20. Yasuhara, K. et al. Spontaneous lipid nanodisc fomation by amphiphilic polymethacrylate copolymers. J. Am. Chem. Soc. 139, 18657–18663 (2017).

21. Workman, C. E. et al. Alternatives to Styrene- and Diisobutylene-Based Copolymers for Membrane Protein Solubilization via Nanodisc Formation. Angew. Chem. Int. Ed 62, e202306572 (2023).

22. Kopf, A. H. et al. Synthesis and evaluation of a library of alternating amphipathic copolymers to solubilize and study membrane proteins. Biomacromolecules (2022) doi:10.1021/acs.biomac.1c01166.

23. Walker, G. et al. Oligomeric organization of membrane proteins from native membranes at nanoscale spatial and single-molecule resolution. Nat. Nanotechnol. 19, 85–94 (2024).

24. Kopf, A. H. et al. Factors influencing the solubilization of membrane proteins from Escherichia coli membranes by styrene-maleic acid copolymers. Biochim. Biophys. Acta Biomembr. 1862, 183125 (2020).

25. Hawkins, O. P. et al. Membrane protein extraction and purification using partially-esterified SMA polymers. Biochim. Biophys. Acta Biomembr. 1863, 183758 (2021).

26. Shi, L. et al. Preparation and characterization of SNARE-containing nanodiscs and direct study of cargo release through fusion pores. Nat. Protoc. 8, 935–948 (2013).

27. Angeletti, C. & Nichols, J. W. Dithionite quenching rate measurement of the inside-outside membrane bilayer distribution of 7-nitrobenz-2-oxa-1,3-diazol-4-yl-labeled phospholipids. Biochemistry 37, 15114–15119 (1998).

28. Kleusch, C., Hersch, N., Hoffmann, B., Merkel, R. & Csiszár, A. Fluorescent lipids: functional parts of fusogenic liposomes and tools for cell membrane labeling and visualization. Molecules 17, 1055–1073 (2012).

29. Cunningham, R. D. et al. Iterative RAFT-Mediated Copolymerization of Styrene and Maleic Anhydride toward Sequence- and Length-Controlled Copolymers and Their Applications for Solubilizing Lipid Membranes. Biomacromolecules 21, 3287–3300 (2020).

30. Whitelegge, J. P., Gundersen, C. B. & Faull, K. F. Electrospray-ionization mass spectrometry of intact intrinsic membrane proteins. Protein Sci. 7, 1423–1430 (1998).

31. van Gestel, R. A. et al. Quantitative erythrocyte membrane proteome analysis with Blue-native/SDS PAGE. J. Proteomics 73, 456–465 (2010).

32. Lujan, P. et al. Sorting of secretory proteins at the trans-Golgi network by TGN46. (2023) doi:10.7554/eLife.91708.1.

33. Stenius, K., Janz, R., Südhof, T. C. & Jahn, R. Structure of synaptogyrin (p29) defines novel synaptic vesicle protein. J. Cell Biol. 131, 1801–1809 (1995).

34. Bera, M. et al. Synaptophysin chaperones the assembly of 12 SNAREpins under each ready-release vesicle. Proc Natl Acad Sci USA 120, e2311484120 (2023).

35. Adams, D. J., Arthur, C. P. & Stowell, M. H. B. Architecture of the synaptophysin/synaptobrevin complex: structural evidence for an entropic clustering function at the synapse. Sci. Rep. 5, 13659 (2015).

36. Olivas, T. J. et al. ATG9 vesicles are incorporated into nascent autophagosome membranes. FASEB J. 36 **Suppl 1**, (2022).

37. Panda, A. et al. Direct determination of oligomeric organization of integral membrane proteins and lipids from intact customizable bilayer. Nat. Methods 20, 891–897 (2023).

38. Panda, A., Brown, C. & Gupta, K. Studying Membrane Protein-Lipid Specificity through Direct Native Mass Spectrometric Analysis from Tunable Proteoliposomes. J. Am. Soc. Mass Spectrom. 34, 1917–1927 (2023).

39. Salem, M., Bernach, M., Bajdzienko, K. & Giavalisco, P. A Simple Fractionated Extraction Method for the Comprehensive Analysis of Metabolites, Lipids, and Proteins from a Single Sample. J. Vis. Exp. (2017) doi:10.3791/55802.

40. Cox, J. et al. Accurate proteome-wide label-free quantification by delayed normalization and maximal peptide ratio extraction, termed MaxLFQ. Mol. Cell. Proteomics 13, 2513–2526 (2014).

