## Supplementary material for "A proteome-wide quantitative platform for nanoscale spatially resolved extraction of membrane proteins into native nanodiscs": SI Table

**Supplementary Information Table 1: Structures solved using membrane active polymers**

| Protein | Polymer | Year | Purified From | Resolution | Reference |
|---|---|---|---|---|---|
| ACRB | SMA 2:1 | 2018 | E. coli | 8.8 Å | [40] |
| ACIII | SMA 3:1 | 2018 | F. johnsoniae | 3.4 Å | [15] |
| ZipA:FtsZ | SMA 2:1 | 2019 | E. coli | 16 Å | [41] |
| KimA | SMA 2:1 | 2020 | E. coli | 3.7 Å | [42] |
| hTRPM4 | AASTY-B | 2020 | HEK293 | 18 Å | [18] |
| BdSLAC1 | SMA (not specified which type) | 2021 | S. pombe | 2.97 Å | [43] |
| Cyt $bo_3$ | SMA 3:1 | 2021 | E. coli | 2.55 Å | [44] |
| ELIC | SMA 3:1 | 2021 | E. coli | 2.5 Å | [45] |
| Yna1 | SMA 2:1 | 2021 | E. coli | 2.4 Å | [46] |
| GlyR | SMA2:1 | 2021 | Sf9 | 3.2 Å | [47] |
| Bam complex | SMA 2:1 | 2022 | E. coli | 3.6 Å | [48] |
| cASIC1 | SMA 2:1 | 2020 | HEK293S GnTI | 2.8 Å | [49] |
| HIV-1 Env | SMA 2:1 | 2023 | A549 | 4.1 Å | [50] |
| WbaP | SMA 2:1 | 2023 | E. coli | Not specified | [51] |
| Cyt $bc_1$ | SMA (not specified which type) | 2023 | R. sphaeroides | 2.9 Å | [52] |
| GPIb-1X-V | SMA 3:1 | 2023 | Expi293F | 11 Å | [53] |

**Supplementary Information Table 2: Polymer description**

| Polymer Name | Description | Synthesis Methods | Origin |
|---|---|---|---|
| SMA140 | Styrene, Maleic Acid ratio 1:1; MW 6.5 KDa | CSTR | Commercial |
| SMA200 | Styrene, Maleic Acid ratio 2:1; MW 6.5 KDa | CSTR | Commercial |
| SMA300 | Styrene, Maleic Acid ratio 3:1; MW 6.5 KDa | CSTR | Commercial |
| AASTY645 | Styrene, Acrylic Acid ratio 55:45; MW 6 KDa | RAFT | Commercial |
| AASTY650 | Styrene, Acrylic Acid ratio 50:50; MW 6 KDa | RAFT | Commercial |
| AASTY655 | Styrene, Acrylic Acid ratio 45:55; MW 6 KDa | RAFT | Commercial |
| AASTY1145 | Styrene, Acrylic Acid ratio 55:45; MW 11 KDa | RAFT | Commercial |
| AASTY1150 | Styrene, Acrylic Acid ratio 50:50; MW 11 KDa | RAFT | Commercial |
| AASTY1155 | Styrene, Acrylic Acid ratio 45:55; MW 11 KDa | RAFT | Commercial |
| CS20 | Chloro Styrene, Maleic Acid ratio 1:1; MW 6.0 KDa | RAFT | Home made |
| CS40 | Chloro Styrene, Maleic Acid ratio 1:1; MW 6.6 KDa | RAFT | Home made |
| CS60 | Chloro Styrene, Maleic Acid ratio 1:1; MW 7.7 KDa | RAFT | Home made |
| CS80 | Chloro Styrene, Maleic Acid ratio 1:1; MW 8.8 KDa | RAFT | Home made |
| AASTY80 | Styrene, Acrylic Acid ratio 55:45; MW 5.1 KDa | RAFT | Home made |

**Supplementary Information Table 3: Extraction conditions**

| Extraction Condition | Buffer Condition | Salt Concentration | 10% glycerol | % polymer |
|---|---|---|---|---|
| SMA140 | 20 mM HEPES pH 7.5 | 100mM NaCl | N | 1% |
| SMA200 | 50 mM TrisHCl pH 8.1 | 300mM NaCl | Y | 1% |
| SMA300 | 20 mM HEPES pH 7.5 | 100mM NaCl | N | 1% |
| AASTY645 | 20 mM HEPES pH 7.5 | 100mM NaCl | N | 1% |
| AASTY650 | 20 mM HEPES pH 7.5 | 100mM NaCl | N | 1% |
| AASTY655 | 20 mM HEPES pH 7.5 | 100mM NaCl | N | 1% |
| AASTY1145 | 20 mM HEPES pH 7.5 | 100mM NaCl | N | 1% |
| AASTY1150 | 20 mM HEPES pH 7.5 | 100mM NaCl | N | 1% |
| AASTY1155 | 20 mM HEPES pH 7.5 | 100mM NaCl | N | 1% |
| CS20 | 50 mM TrisHCl pH 8.1 | 150 mM NaCl | Y | 1.5% |
| CS40 | 50 mM TrisHCl pH 8.1 | 150 mM NaCl | Y | 1.5% |
| CS60 | 50 mM TrisHCl pH 8.1 | 150 mM NaCl | Y | 1.5% |
| CS80 | 50 mM TrisHCl pH 8.1 | 150 mM NaCl | Y | 1.5% |
| AASTY80 | 50 mM TrisHCl pH 7.8 | 150 mM NaCl | Y | 1.5% |
